# An Infusible Extracellular Matrix Biomaterial Improves Survival in a Model of Severe Systemic Inflammation

**DOI:** 10.1101/2024.06.05.597616

**Authors:** Maria Karkanitsa, Raymond M. Wang, Anne C. Lyons, Joshua M. Mesfin, Alexander Chen, Martin T. Spang, Filiberto Quintero, Kaitlyn Sadtler, Mark Hepokoski, Karen L. Christman

**Author notes:** These authors contributed equally to this work (Raymond M. Wang, Anne C. Lyons). **Corresponding Author:** Karen L. Christman, PhD, Sanford Consortium for Regenerative Medicine 2880 Torrey Pines Scenic Drive, La Jolla, CA 92037. **Author Contributions:** MK designed the study, performed experiments, analyzed data, interpreted results, and drafted the manuscript. RW and AL designed the study, performed experiments, analyzed data, and interpreted results. JM, AC, MS, and FQ performed experiments and analyzed data. MH designed the study and interpreted the results. KLC designed the study, interpreted results, and obtained funding. KS aided in design of the flow cytometry study. All authors reviewed and edited the manuscript and approved the final version.

## Abstract

Excess systemic inflammation can often be lethal in septic and trauma patients due to onset of multiple organ dysfunction syndrome (MODS). As of right now, there are no effective immunomodulatory therapeutics that can promote survival within this patient population. Pro-regenerative extracellular matrix (ECM) biomaterials have shown success for treatment of local inflammation but have not been fully explored for treating systemic inflammation. Here, we demonstrate efficacy of an intravenously delivered infusible ECM (iECM) material, which promotes increased survival in a murine model of MODS by decreasing systemic mediators of inflammation. Lung and kidney failure are associated with higher mortality in MODS compared to other organ failures, and we demonstrate that iECM localizes primarily to kidney and lung tissues during systemic inflammation induced by endotoxin. iECM successfully lowered vascular permeability within lung tissue and lowered levels of inflammatory cytokine signaling such as IL-6, verified via ELISA and gene expression analyses. We also demonstrated that immune cell infiltration into lung tissue was modulated with iECM treatment, with an increase in neutrophil retention in the lung and decreases in pro-inflammatory macrophage presence. In summation, iECM improves survival from severe systemic inflammation by decreasing the local and systemic inflammatory signaling pathways that contribute to MODS. These results provide a strong rationale for translational studies of iECM treatment in systemic inflammatory syndromes, including sepsis and trauma.

## INTRODUCTION

While inflammation is necessary for effective clearance of pathogens, excess systemic immune activation can lead to tissue damage and improper organ function. Multiple organ failure dysfunction syndrome (MODS) is a well-characterized complication in trauma and septic patients that is characterized by dysfunction of two or more organ systems, mainly due to aberrant inflammation ^1,2^. While timely administration of antimicrobials, intravenous fluids, and vasopressors has improved outcomes, the mortality rate for patients suffering from MODS remains unacceptably high at 25-35% ^1^.

While anti-inflammatory treatments for MODS have shown benefit in preclinical models ^3–5^, there still remains a clinical need for better immunomodulatory therapeutics within this patient population. Furthermore, several preclinical studies for immunomodulatory therapeutics to treat severe systemic inflammation have only shown efficacy when therapeutic intervention occurs prior to induction of an infectious or inflammatory insult ^5–7^. Unfortunately, this approach does not accurately reflect the clinical timeline of most critically ill patients, who present to the hospital once systemic inflammation has begun. As a result, no proven pharmacological treatments for most systemic inflammatory syndromes, including sepsis and trauma, exist. To date, pro-regenerative, anti-inflammatory biomaterial strategies have not been explored for this indication.

Recently, we developed an infusible decellularized extracellular matrix (iECM) that can be delivered intravascularly. This novel iECM binds to leaky inflamed vasculature, has pro-regenerative and immunomodulatory effects, and reduces microvascular permeability in preclinical MI models ^8^. Microvascular leak not only impacts tissue perfusion, but also promotes infiltration of inflammatory cells that contribute to MODS^9^. We therefore hypothesized that iECM would be an effective treatment strategy for MODs via reduction in vascular permeability and promotion of an anti-inflammatory immune response. To test this hypothesis, we evaluated intravenous delivery of iECM compared to saline in a mouse lipopolysaccharide (LPS) model, which is a commonly used model for sepsis, acute respiratory distress syndrome (ARDS), and other multiple organ dysfunction syndromes.

## RESULTS

### iECM Targets Sites of Inflammation and Lowers Microvascular Leakage Within Lung Tissue

To model the systemic inflammatory effects of sepsis, we utilized a dual dosage system of LPS (**Figure 1a**) ^10,11^. It is hypothesized that full organ dysfunction due to aberrant inflammation is likely due to a minimum of two separate stimuli, termed the ‘two-hit’ model ^12^. We found that a dual-dose LPS system was able to effectively dysregulate homeostasis as verified by drop in core body temperature (**Supplemental Figure 1A**) and body weight (**Supplemental Figure 1B**) and cause high mortality rates over a 30 hour period, mimicking the acute mortality rates reported in severe septic patients with severe organ failure ^13^.

**Figure 1.**
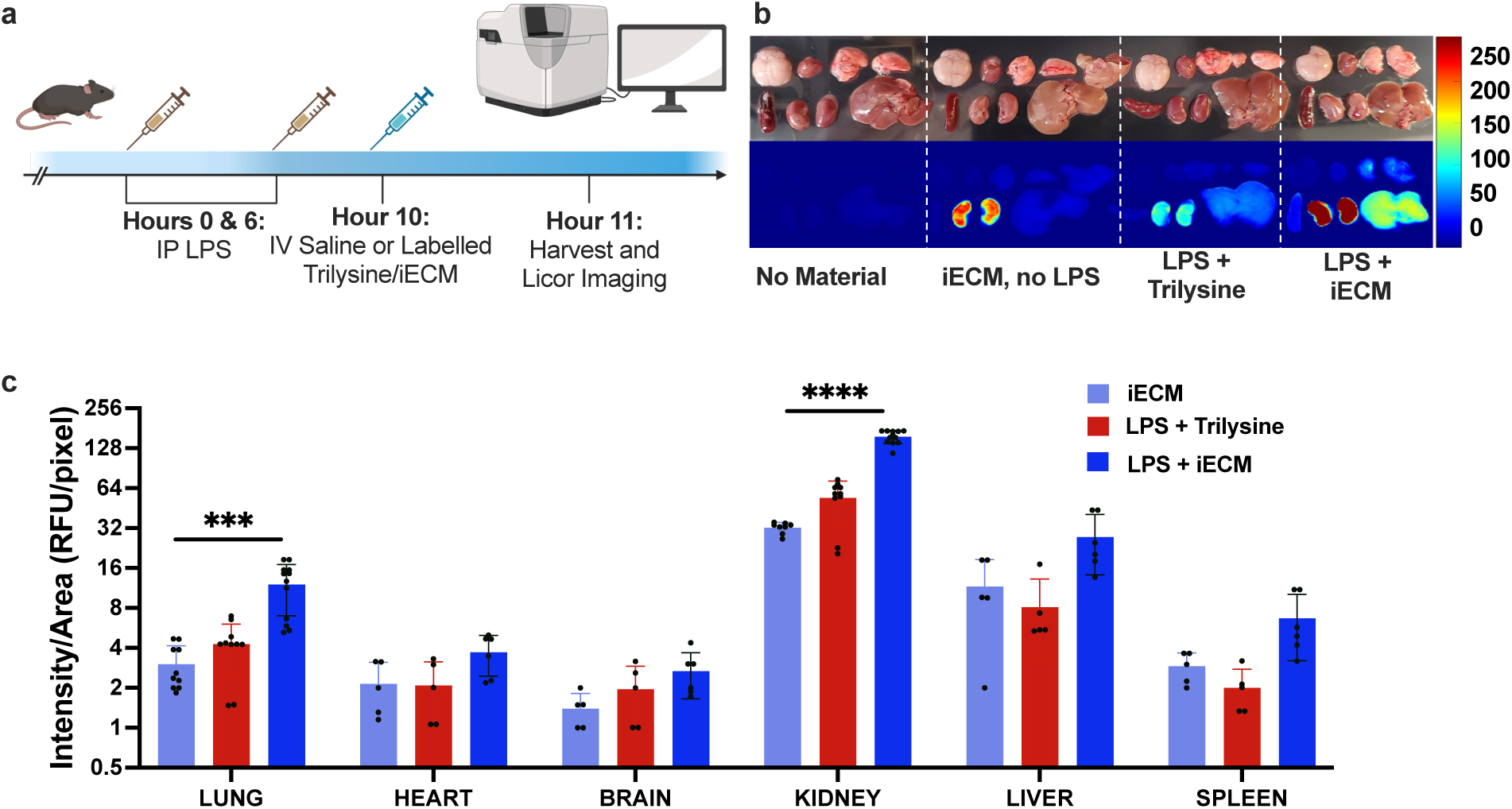
Biodistribution of iECM in Animal Model of MODS. a. Overview of timeline for biodistribution assessment in healthy animals with no treatment, healthy animals treated with iECM, LPS-dosed animals receiving an IV injection of trilysine, and LPS-dosed animals receiving an IV injection of iECM. b. Brightfield and Licor imaging at 785 nm to visualize vivotag-750 tagged iECM and trilysine. c. Signal intensity of vivotag-750 tagged iECM and trilysine were quantified and compared across each organ. Significance was determined using an unpaired t-test assuming individual variance for each organ. P values were adjusted using the Holm-Šídák method. *p < 0.05, **p < 0.01, ***p < 0.001, and ****p < 0.0001.

Once our model was established, we sought to assess retention and biodistribution of iECM in LPS-treated animals. While we previously identified sites of leaky vasculature as sites of iECM retention in local injury and disease models ^8^, iECM retention has not been previously evaluated in a model of systemic inflammation. We noted that iECM retention was increased across all tissues after LPS administration compared to non-LPS treated animals (**Figure 1b**). To determine if this was due to specific binding or by passive diffusion through leaky vasculature due to a phenomenon known as the enhanced permeability and retention effect, we compared retention of our iECM to a trilysine peptide control. Relative to trilysine, we observed that iECM accumulation was higher in the lungs and kidney (**Figure 1c**). This suggests there is an active mechanism of iECM binding and retention within these tissues.

To evaluate how iECM affects microvascular permeability within this model of MODS, we treated animals with iECM or saline and intravenously administered fluorescently labeled albumin as a metric for tissue perfusion (**Figure 2a**, **Supplemental Figure 2**). We evaluated multiple organs; however, the lung was of particular interest given that edema within lung tissue is associated with high mortality rates in critical care patients and is a hallmark of respiratory failure and ARDS ^14,15^. We determined that iECM significantly limited leakage of fluorescently labelled albumin within lung tissue, confirming that iECM lowers vascular permeability in lung tissues (**Figure 2b and 2c, Supplemental Figure 2**). Albumin leakage was not significantly altered in other tissues (**Supplemental Figure 3**).

**Figure 2.**
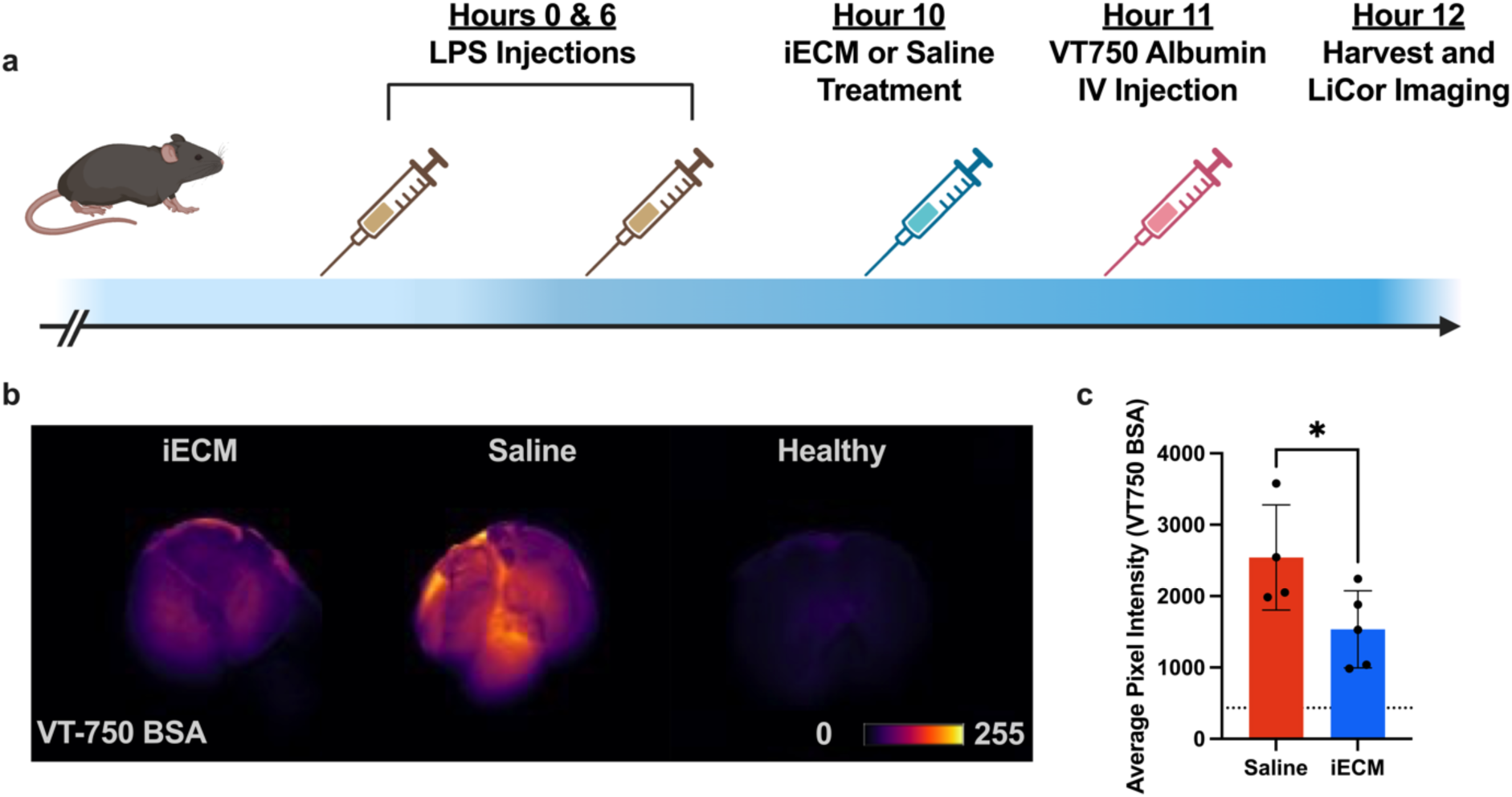
Vessel permeability is lowered in the lung with iECM treatment. a. Overview of the animal timeline used to quantify vessel permeability. b. Representative images of VT-750 albumin of lungs from animals treated with iECM or saline compared to healthy control lungs injected with albumin. c. Quantification of fluorescently labelled albumin in lungs of saline (red) and iECM (blue) treated animals. Statistical significance was determined using a two-tailed, unpaired t-test. *p <0.05.

### iECM Promotes Survival Post Severe Inflammatory Insult

We further characterized the therapeutic effects of the iECM in this endotoxin model by tracking animal survival over a 30 hour period as depicted in **Figure 3a**. Compared to the saline-treatment control, iECM significantly improved animal survival with an average increase of 27.4% at 30 hours (**Figure 3b**). At 30 hours, tissues were processed and evaluated at the proteomic, transcriptomic, and cellular levels for differences due to iECM treatment (**Figure 3c**).

**Figure 3:**
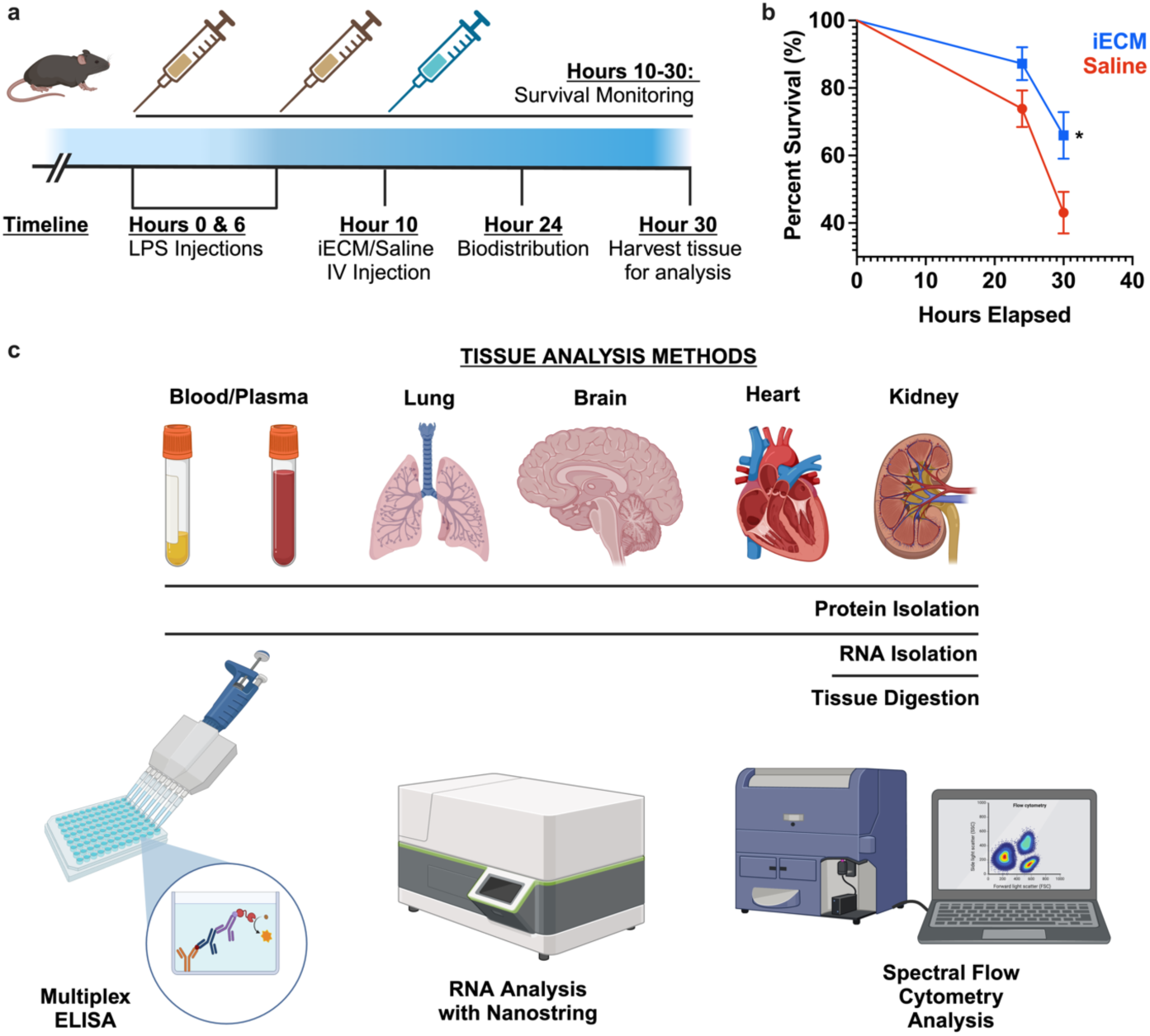
Overview of methods used to determine iECM efficacy in MODS model. a. Overview of animal model timeline. Animals were treated with a dual-dose of LPS as described previously and then treated with saline or iECM. Survival was monitored over a 30-hour time period. At 30 hours, animals that survived were harvested for analysis. b. A total of n = 112 animals (n = 47 for iECM, n = 65 for saline) were utilized for analysis and incorporated for survival analysis. Survival curve analysis was performed using the Log-Rank (Mantel Cox) test. c. In addition to survival analysis, blood/plasma, brain, heart, kidney, and lung tissue was harvested and processed for protein analysis using a multikine ELISA system and transcriptomic analysis using the Nanostring nCounter platform. Lung tissue was further processed for a 22-color spectral flow cytometry panel. Error bars= standard error of the mean.

### Treatment with iECM Decreases Cytokine Expression Systemically

Building on our hypothesis that iECM would promote survival via immunomodulation, we sought to further characterize how the iECM altered immune signaling after LPS administration. To do this, we assessed changes in cytokine levels due to iECM treatment using a multiplex ELISA system. Cytokines are well-established contributors to MODS, with rising levels of Interleukin 6 (IL-6) and tumor necrosis alpha (TNF-α) correlating to MODS progression in clinical studies ^16–18^. We probed a total of 13 cytokines in tissue lysates and plasma to determine both local and systemic immune states. In solid tissues, we saw downregulation of IL-6, IFNγ, IL-17A, and IL1b by iECM. We noted that, of the 13 cytokines screened, IL-6 had the largest decrease and was significantly lowered in the lung, heart, kidney, and plasma of iECM-treated animals (**Figure 4a**). IL-6 levels also trended downward in the spleen (p = 0.08) and the brain (p = 0.19). We also noted decreased interferon gamma levels in the kidney of iECM-treated animals but not in other tissues (**Figure 4b**). IL-1α was additionally downregulated in the lung, and IL-1β was lowered in the spleen of iECM-treated animals (**Figure 4c and 4d**) while IL-17A trended downwards in the spleen (**Figure 4h**; p = 0.06). Compared to the solid tissues probed, we observed that the differences in cytokine levels were most pronounced in plasma. IL-6 (**Figure 4a**), IL-1α (**Figure 4c**), IL-1β (**Figure 4d**), GM-CSF (**Figure 4e**), TNF-α (**Figure 4f**), IL-12p70 (**Figure 4g**), and IL-23 (**Figure 4i**) were all significantly downregulated in the plasma. Levels of plasma IL-10 trended downwards with iECM but remained nonsignificant along with levels of MCP-1, IL-27, and IFN-β (**Supplemental Figure 4**).

**Figure 4.**
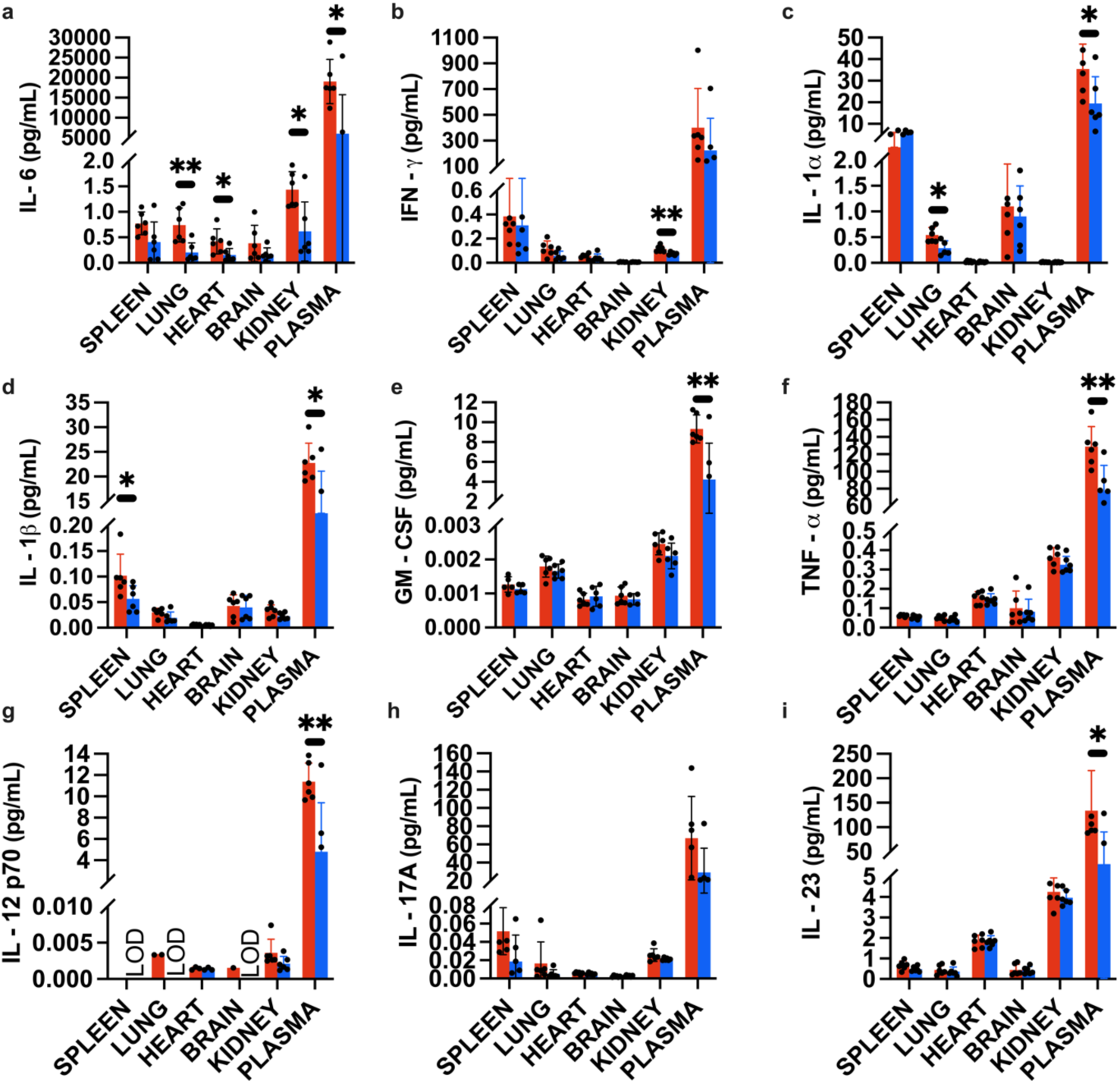
Quantification of inflammatory cytokines. Cytokine levels were assessed in plasma and tissue lysate from saline (red, n = 6) and iECM (blue, n = 6) treated animals using a 13-plex multiplex ELISA array. A. IL-6, B. IFN-y, C. IL-1, D. IL-1b, E. GM-CSF, F. TNF-a, g. IL-12 p70, H. IL-17, I. IL-23. Significance was determined using an unpaired t-test with individual variance assumption for each row to accommodate for inherent differences in baseline tissue expression of each cytokine. Data is mean ± standard deviation. *p < 0.05, **p < 0.01. LOD: Limit of Detection, indicating levels fell below detectable levels of the assay.

### iECM Treatment is Associated with Transcriptional Downregulation of Pro-Inflammatory Pathways

We further evaluated immunomodulatory mechanisms by assessing transcriptional differences across all tissues via Nanostring nCounter gene expression analysis. mRNA was isolated from whole blood, lung, brain, heart, and kidney tissues and transcriptional differences were assessed using a panel of 547 genes involved in the immune response. Analysis of probed tissues revealed that the greatest number of differentially expressed genes were present in the blood (**Supplemental Table 1**), followed by lung (**Supplemental Table 2**), brain (**Supplemental Table 3**), heart (**Supplemental Table 4**), and kidney (**Supplemental Table 5**). In the blood, transcriptional downregulation of *Csf2*, *Il6*, and *Il1a* matched the ELISA data showing lower levels of these cytokines in blood (**Figure 5a**). Additionally, we noted that genes encoding for chemokines including *Ccl11*, *Ccl2*, and *Cxcl15* were downregulated in blood. Chemokines are a subset of cytokines involved in cellular motility and recruitment during inflammation, and their unregulated release has been associated with negative outcomes in clinical trials of severe trauma ^19–21^. We also observed decreased gene expression of adhesion molecules in blood with iECM treatment, such as VCAM1, ICAM1, and NCAM1 (**Supplemental Table 1**).

**Figure 5.**
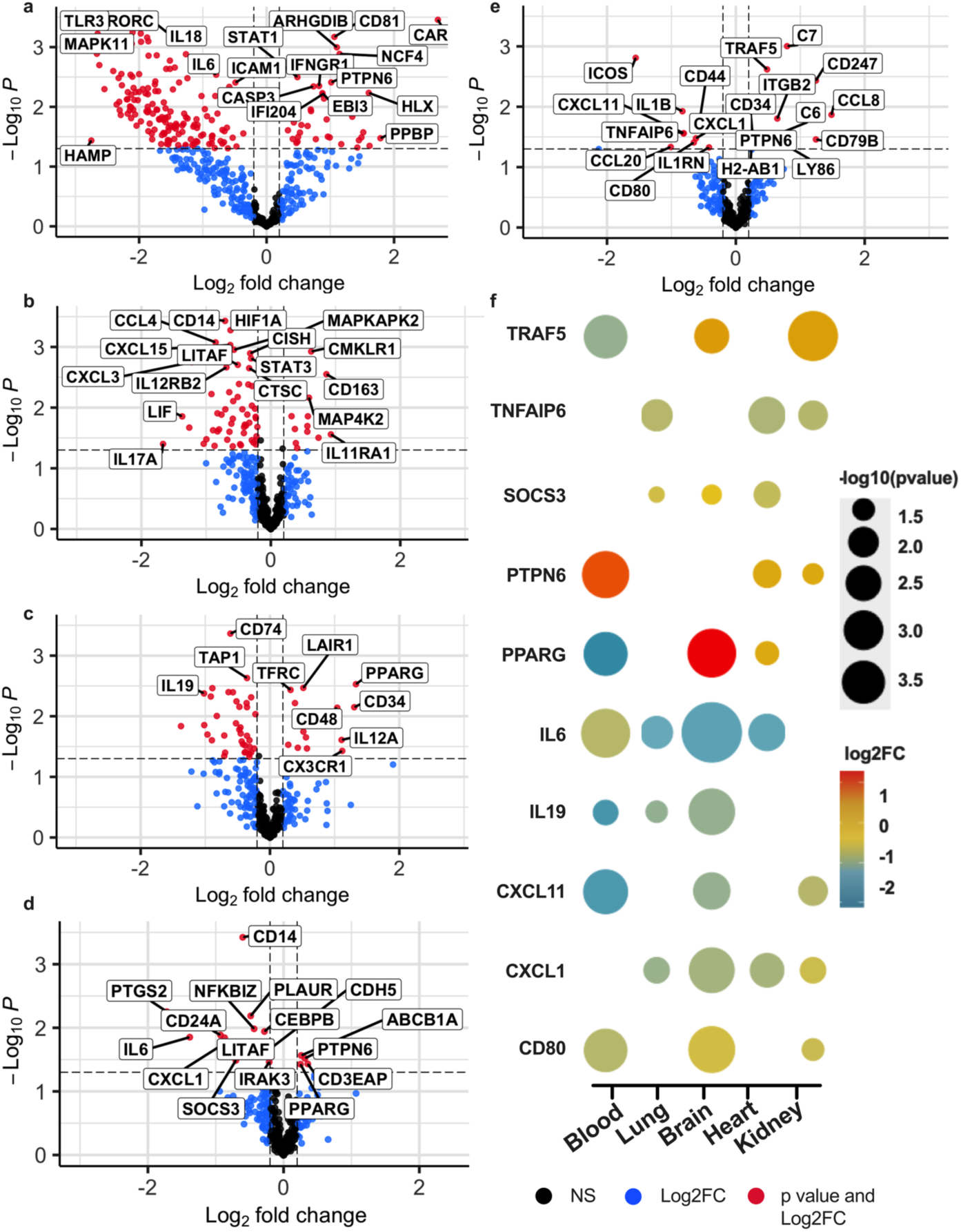
Volcano plot of differentially expressed genes in iECM treated animals versus saline. Nanostring gene expression analysis was performed on samples from A. blood, B. lung, C. brain, D. heart and E. kidney. An n = 5-6 were used per tissue. Nanostring analysis was performed using Rosalind to determine log fold change and p values for each gene. Differentially expressed genes that were significant (p < 0.05) and had an absolute fold change >1.01 are colored red, while non-significant genes that were differentially expressed are shown in blue. F. Genes that were commonly differentially expressed (> 3 tissues) were plotted together to show trends in downregulated genes. Size of dots correlate to −log10(pvalue) in gene expression, while color correlates to the log2 fold change in gene expression in iECM versus saline-treated animals.

Given that intraperitoneal injection of LPS has been previously used to study ARDS ^10,14^, we were particularly interested in the immunological differences in the lungs of iECM-treated animals. There were several notable genes within the lung that were differentially expressed (**Figure 5b**). Similar to the ELISA data, we observed downregulation of genes encoding Il-α (*Il1a)* and IL-6 (*Il6*), aligning with decreased levels of these cytokines. We also noted decreased gene expression of several chemokines as we saw in the blood, including *Ccl3*, *Cxcl1*, *Ccl5*, and *Ccl4*, suggesting altered immune cell recruitment to the lung. Particularly relevant in the lung was observed downregulation of *Cxcl15*, known as lungkine or IL-8. IL-8 has been previously used as a biomarker for ARDS and has been negatively correlated with survival in clinical studies ^17,22^. Its downregulation in the lung suggests that the iECM lowers lung damage in this MODS model. We noted that, in addition to downregulation of inflammatory mediators, *Hif1a*, a gene encoding for Hypoxia-Induced Factor 1 alpha (HIF1α) was significantly lower in iECM-treated animals. This suggests that hypoxia and hypoxia-induced inflammatory signaling could be minimized as a result of iECM treatment, possibly due to lowering of hyperinflammation-induced tissue damage. In addition, we noted that one of the upregulated genes within the lung was *Cd163*, which is a known marker for M2 ‘alternatively activated’ macrophages.

Encephalopathy due to hyperinflammatory signaling is well-established in clinical studies of cytokine storm ^23^, and neuroinflammation is exacerbated by increased levels of inflammatory chemokines such as *Ccl5*, aka RANTES ^24^. Within the brain, we found that the majority of notable downregulated genes were pro-inflammatory chemokines such as *Cxcl11, Ccl2, Ccl5,* and *Cxcl1* (**Figure 5c**). We also observed downregulation of *Fas* and *Casp8*, both of which are involved in FAS-signaling that is well established to induce apoptotic cell death ^25^.

In the heart, transcription of inflammatory cytokine *Il6* and chemokine *Cxcl1* were both downregulated (**Figure 5d**). Interestingly, *Ptgs2,* the gene encoding for Prostaglandin 2 or COX-2, was the gene downregulated with the greatest magnitude,. COX-2 is a main target of common non-steroidal anti-inflammatory drugs (NSAIDs), and its therapeutic targeting has been shown to improve cardiac dysfunction in previous murine models of LPS-induced MODS ^26^.

Within the kidney, inflammatory chemokines *Cxcl1, Cxcl11*, and *Ccl20* were downregulated in iECM treated animals (**Figure 5e**). *Icos*, a known marker of stimulated T cells, was also significantly downregulated ^27,28^. While the decrease in IL-1β protein levels was only trending in the ELISA data (**Figure 4d**), *Il1b* was significantly downregulated in Nanostring analysis (**Figure 5e**).

We observed that several inflammatory cytokines and chemokines were consistently downregulated across several tissues (**Figure 5f**). Several differentially expressed genes in iECM-treated tissues were ones involved in signal transduction, including *Traf5*, *Tnfaip6*, *Socs3*, *Ptpn6*, and *Pparg* (**Figure 5f**). Many of these act downstream of cytokine signaling pathways such as TNF-α and IL-6 ^29,30^, suggesting again that these pathways are altered as a result of iECM treatment. Similar to the ELISA results, genes encoding for IL-6 were downregulated in blood, brain, heart, and lung of iECM-treated animals. The gene encoding for proinflammatory chemokine CXCL1 was also significantly downregulated in 4 of the 5 tissues probed (brain, heart, kidney, and lung). *Cxcl11*, another proinflammatory chemokine, was downregulated in the blood, brain, and kidney.

From the number of conserved downregulated genes as well as the downregulation of multiple different cytokines/chemokines across tissues, we utilized the enrichR package in R to perform pathway enrichment analysis using the mouse WikiPathways 2019 library to assess downregulated gene pathways ^31^. PathfindR enrichment analysis was also performed and detailed in **Supplementary Figures 5 – 14**. Pathway enrichment showed that iECM treatment dampened several key inflammatory signals across all tissues. In the blood, IL-6 signaling as well as TNF-alpha signaling was downregulated (**Figure 6a**). In addition to other inflammatory signaling pathways, we observed downregulation of coagulation and complement pathways in blood. In the lung, the top downregulated pathway was TNF-alpha signaling via NF-kb (**Figure 6b**). We also noted decreases in interferon gamma signaling, IL-2 signaling, and apoptotic pathways. In the brain, we noted that several downregulated genes were involved in interferon gamma signaling, TNF-alpha signaling, and IL-6 signaling (**Figure 6c**). In addition, we found downregulation of complement and apoptosis within the brain of iECM-treated animals. In the heart we observed highest downregulation of TNF-alpha signaling (**Figure 6d**) followed by IL-6 and Interferon gamma signaling. There was also reduced hypoxia signaling within the heart tissue that was similarly noted in the kidney tissue (**Figure 6e**). In the kidney, there was also reduction in TNF-alpha signaling, IL-6 signaling, and the interferon gamma response. Apart from interferon alpha response in the heart, there were 9 immune signaling pathways downregulated across all five tissues probed, providing further support that iECM lowers systemic levels of hyperinflammation (**Figure 6f**).

**Figure 6.**
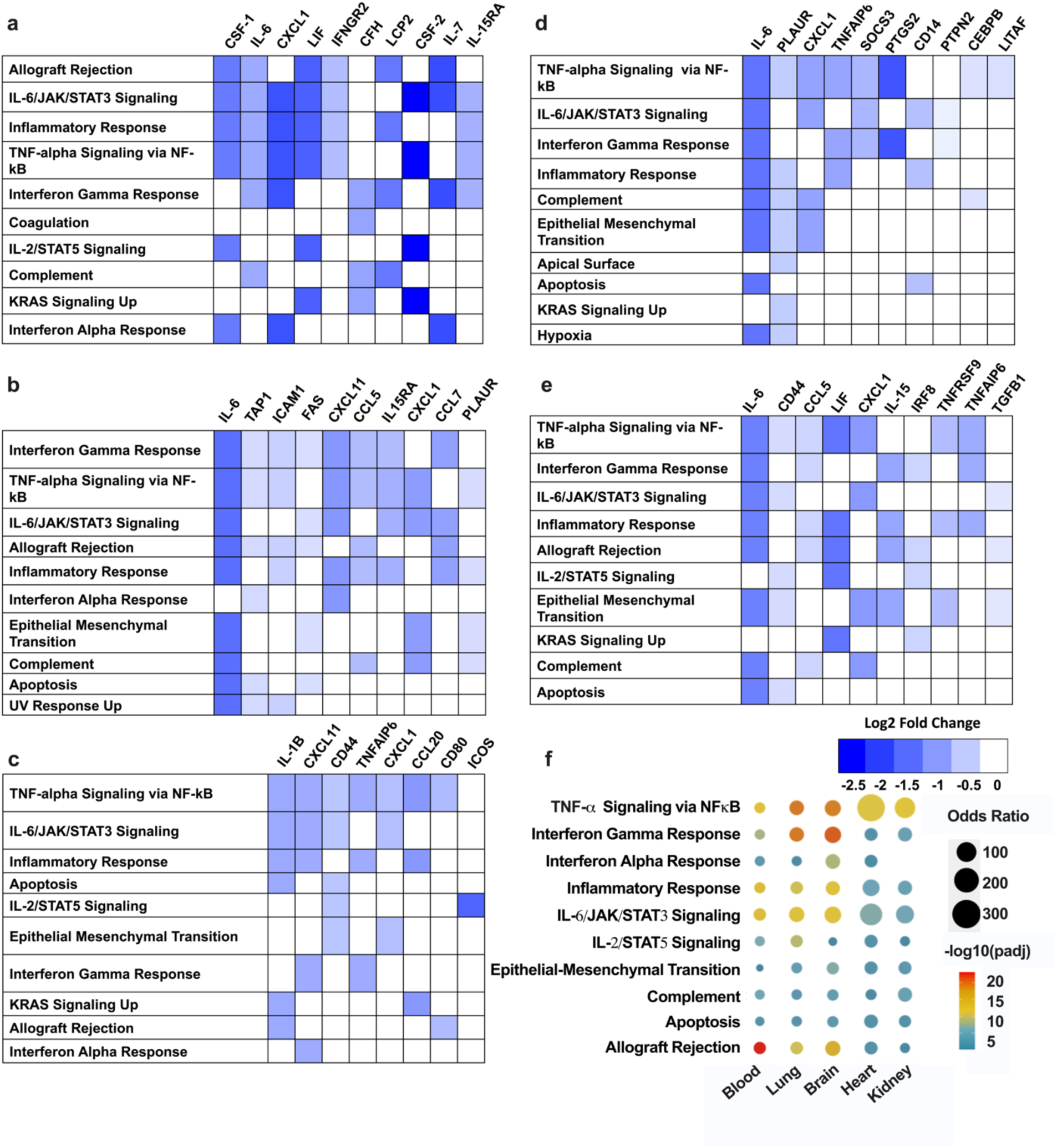
Downregulated immune signaling pathways in tissues of iECM-treated animals compared to saline. Enrichment analysis was performed using EnrichR to determine significantly downregulated immune signaling pathways from downregulated genes in each tissue. Genes as well as their log2 fold change in common pathways were incorporated to show crosstalk across immune signaling. A. Blood, B. Lung, C. Brain, D. Heart, E. Kidney. F. To identify commonly downregulated pathways, the top 10 significantly differentially expressed signaling pathways in all tissues were compared. P values were adjusted using the Benjamini-Hochberg method. Color = −log10(padj) and dot size = Odds Ratio.

### Flow Cytometry Demonstrates Decreased Presence of Several Inflammatory Cell Types with iECM Treatment

To further assess immune cell differences arising due to iECM treatment, we utilized a previously established 22 color spectral flow cytometry panel^32^ developed for assessing myeloid cell responses to implanted biomaterials (panel outlined in **Supplemental Table 6**). The gating strategy utilized to identify immune cell populations can be seen in **Supplementary Figure 15** with full FlowSOM characterization of CD45+ immune cells shown in **Supplementary Figures 16-21**. Macrophage phenotypes were identified by the gating shown in **Supplementary Figure 22**. We initially performed UMAP clustering of CD45+ cells in lungs of saline and iECM-treated animals (**Figure 7a**) and observed differences in the UMAP-established clusters. Upon looking at immune cell populations using the gating strategy outlined in **Supplemental Figure 15**, we confirmed that our cell populations matched UMAP-derived clusters of cells well (**Figure 7b**). Upon comparing cell populations in saline and iECM-treated animals, it was clear that neutrophil levels were significantly increased in the lung with iECM treatment (**Figure 7c**) while levels of CD11b-lymphoid cells trended lower with iECM treatment. While our flow cytometry panel was designed to look at myeloid cell populations, FlowSOM clustering revealed two separate clusters of CD11b-MHCII-cell populations (**Figure 7d**). Using ClusterProfiler, we were able to identify this cell population as a population of Ly6C+ lymphoid cells (**Figure 7e and 7f**). Ly6C expression has been reported on various immune cells such as natural killer and T cells ^33–35^. Utilizing gating of our CD45+ cells, we identified that Ly6C+ CD11b-MHCII-immune cells were significantly lower in the iECM (**Figure 7g**).

**Figure 7.**
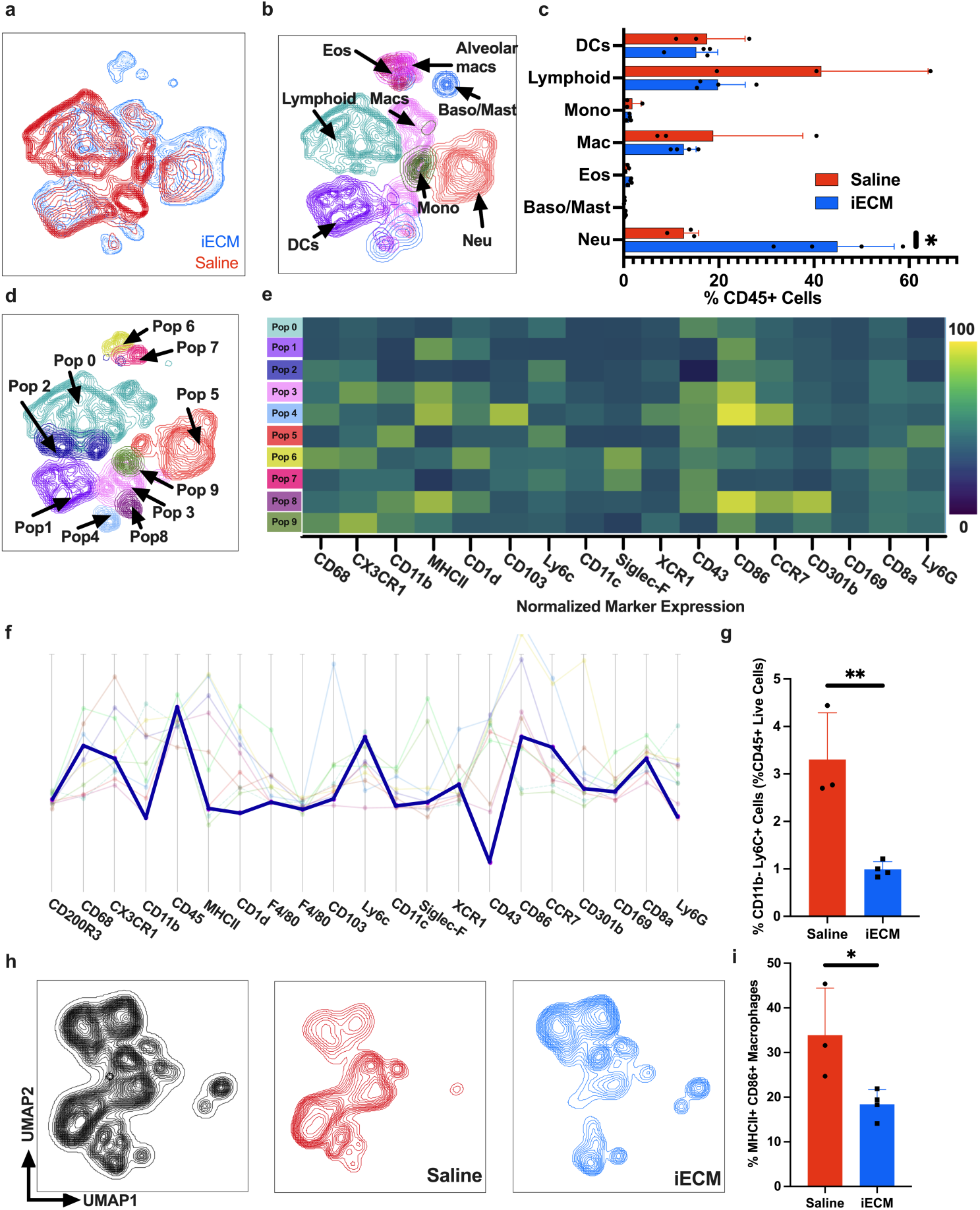
Flow Cytometry Analysis of immune cells in the lung of animals treated with iECM or saline. A. UMAP of live CD45+ Cells isolated from saline (red) and iECM (blue) animals B. UMAP showing cells classified by gating strategy. C. Percentages of major cell populations in lung of saline vs iECM treated animals. D. Unsupervised clustering of all CD45+ cells using flowSOM analysis. E. Normalized marker expression in FlowSOM-derived immune cell clusters. F. Relative marker expression of CD45+ immune cells in cluster 2. G. Comparison of levels of gated CD11b-Ly6C+ cells in lungs of saline (red) vs iECM (blue) treated animals based on relative expression of markers in cluster two of FlowSOM-derived clusters. H. UMAP of macrophages as determined using the gating strategy outlined in Supplemental Figure 8. I. Percentages of inflammatory macrophage phenotypes in saline vs. iECM-treated animals based on expression of inflammatory markers MHCII and CD86. Significance was determined using a two-tailed unpaired t-test. n = 4 per groups. *p < 0.05.

We next looked at the macrophage compartment to characterize differences in macrophage phenotype with iECM treatment. To do this, we utilized the gating strategy outlined in **Supplementary Figure 15** to isolate macrophages as well as the gating strategy in **Supplementary Figure 22** to further evaluate macrophage populations. We immediately noticed with UMAP clustering that macrophages clustered differently due to iECM treatment (**Figure 6H, Supplementary Figures 23-25**). We were also able to observe decreased levels of macrophages expressing proinflammatory ‘M1’ markers MHCII and CD86 (**Figure 6H**). Furthermore, there were significantly lower levels of CD1d+ macrophages with iECM treatment (**Supplementary Figure 26**).

## DISCUSSION

We have shown that a pro-regenerative, infusible, decellularized ECM biomaterial can mitigate systemic inflammation and promote survival in a murine MODS model. Of note, the iECM improved survival when delivered 4 hours following a second dose of LPS, which mimics the clinical scenario where a therapeutic would be delivered after systemic inflammation is already occurring in the patient. This contrasts with several preclinical studies, which have only shown improved survival when the therapeutic is delivered prior to introduction of inflammatory stimuli.

ARDS, encephalopathy, acute kidney injury, and cardiomyopathy are all common complications of systemic inflammatory syndromes that increase risk of morbidity and mortality^15,36–38^. Critically, respiratory failure characterized by severe edema results in a reported 30-40% mortality rate among critical care patients^15,39^. We demonstrate that, relative to saline, iECM significantly lowered vascular permeability in the lung, suggesting a reduction in edema. We also show that iECM was immunomodulatory in the lungs, kidneys, brain, and heart, in addition to decreasing circulating inflammatory mediators present in the serum. Specifically, we showed a consistent downregulation of IL-6 signaling, which is a major driver of MODS, with serum IL-6 levels correlating directly with mortality in several inflammatory disease states ^16,40,41^. We also noted that while TNF-α and interferon gamma levels were only lowered in the plasma and kidney, respectively, interferon gamma signaling and TNF-α signaling were downregulated across all 5 tissues probed with Nanostring analysis. This suggests that while protein levels of cytokines at 30 hours were not significantly different, the transcriptional signaling of these pathways are lowered in iECM-treated animals.

We also noted that a large portion of differentially expressed genes were involved in chemokine signaling. ECM degradation products have been previously shown to modulate chemotaxis in vitro, and several publications have since established differential recruitment of pro-remodeling, anti-inflammatory immune cells to sites of ECM scaffold implantation ^42–44^. Chemokine ligands are also upregulated in response to inflammatory stimuli, particularly IL-6 ^40,45^. As a result, we theorize that downregulation of IL-6 and other inflammatory cytokines resulted in lower transcription of genes encoding for chemokines, such as eotaxin, MCP-2, CXCL1, and CXCL15.

In addition to decreased chemokines and cytokines, we also observed downregulation of genes encoding for multiple adhesion proteins in the blood and other tissues, including *Icam1, Ncam1, and Vcam1.* All these adhesion molecules are involved in immune cell infiltration and extravasation and are markers of endothelial cell activation ^46^. We previously shown that iECM increases rat endothelial cell viability and improves metabolic activity in an *in vitro* study ^8^. Lowered expression of cell adhesion molecules suggests that the iECM helps lower endothelial cell activation.

Using a 22-color myeloid flow cytometry panel, we were able to understand alterations in immune infiltration within the lung with iECM treatment. Unexpectedly, we found that a majority of CD45+ immune cells in the lung with iECM treatment were neutrophils. This was surprising, as neutrophils are classically pro-inflammatory and can often contribute to organ dysfunction. However, studies have shown that neutrophils have high heterogeneity and there are a plethora of neutrophil phenotypes, including pro-remodeling phenotypes^47^. Furthermore, neutrophil sequestration in the lung microvasculature has been well-documented, with LPS inducing a neutrophil crawling phenotype that allows for sequestration of bacteria in models of bacteremia without induction of NETosis^48^. As the iECM is also localizing to the microvasculature of the lung to limit vascular leakage, it is possible that the iECM is interacting with these neutrophils and further facilitating their accumulation on the microvascular wall, while also not inducing degranulation. FlowSOM analysis also identified a population of CD11b-MHCII-Ly6C+ lymphoid cells that were significantly lower with iECM treatment. Ly6C is present on a host of cells, including various activated plasma cells and CD4+ T cell subsets ^33,34^. It is possible that these cells contribute to the inflammatory state within the lung and could contribute to decreased survival in saline-treated animals. We also noted decreased levels of several inflammatory macrophage populations within iECM treated animals. Macrophages of saline-treated animals expressed significantly higher levels of M1 markers MHCII and CD86. MHCII and CD86 are all known markers of inflammatory ‘M1’ polarization ^49^. These cells are known to produce inflammatory cytokines including IL-17A, IL-6, and IL-1a, all of which were shown to be transcriptionally downregulated in the lung of iECM-treated animals. We also observed significant downregulation of CD1d expression in macrophages of iECM-treated animals. CD1d presents lipid antigens to invariant natural killer T cells, which have been associated with mediation of cytokine production and inflammation in acute lung injury and following endotoxin treatment ^50,51^. In looking at the macrophages using flowSOM and UMAP analysis, we observed a cluster of macrophages that were seemingly non-existent in iECM-treated animals. Further analysis of these cells showed that they expressed elevated levels of CD68, CX3CR1, MHCII, CD1d, and CD86, all suggesting a proinflammatory phenotype. Downregulation of macrophage activation could also lead to downregulation of inflammatory chemokine production, which we also identified in the lung of iECM-treated animals. The differential immune cell infiltration in the lung as well as alterations in inflammatory macrophage phenotype support the idea that iECM lowers inflammation within the lung and can prevent inflammatory cell recruitment after treatment.

In conclusion, we established proof-of-concept for using iECM as a novel and effective therapeutic in a preclinical MODS model. We demonstrated mitigation of systemic inflammation by assessing cytokine protein levels across several tissues and confirmed these results utilizing both transcriptional and flow cytometry analyses. These results warrant further translational studies and development of the iECM as an immunomodulatory therapeutic for sepsis and other multiple organ failure syndromes.

## Materials and Methods

### Infusible ECM Biomaterial Generation

iECM was generated from porcine left ventricular tissue obtained from adult Yorkshire farm pigs (30-45 kg) based on previously established protocols ^8^. Porcine left ventricular tissue was minced into small pieces and decellularized under mechanical agitation in a solution of phosphate buffered saline (PBS) containing 1% (wt/vol) sodium dodecyl sulfate (SDS) (Fischer Scientific, Fair Lawn, NJ) with 0.5% 10,000 U/mL penicillin streptomycin (PS) (Gibco, Life Technologies, Grand Island, NY) for 4-5 days to fully decellularize based on previously established criteria (1). The decellularized tissue was rinsed for 24 hours with deionized water and then rinsed multiple times in water under high agitation for thorough removal of residual SDS, lyophilized, and milled into a fine powder. The ECM powder was partially digested with 1 mg/mL pepsin in 0.1M HCl solution for 48 hours before solution was neutralized to pH of 7.4 and reconstituted to physiological salt concentrations.

For infusible ECM generation, the insoluble portion of the digested ECM suspension was pelleted by high-speed centrifugation at 15,000 RCF for 45 minutes at 4oC. The suspended ECM supernatant was transferred into Spectra Por Biotech-Grade CE Dialysis Tubing, 100-500 MWCO (Spectrum Chemical, New Brunswick, NJ), and placed in a series of solutions: 0.5x PBS, 0.25x PBS, and 3 times in deionized water for 12-16 hours. Dialyzed ECM was collected, lyophilized, weighed, and resuspended with 1x PBS for a final concentration of 16 mg/mL based on dry weight. A series of Millipore® Stericup® vacuum filtrations through 0.45 μm and 0.22 μm PVDF membranes ensured the generated iECM was sterile. iECM was aliquoted, lyophilized, weighed, and stored at −80°C with desiccate until needed by resuspension in sterile deionized water for a final concentration of 10 mg/ml^8^. iECM was characterized and passed release criteria using sGAG quantification, dsDNA quantification, PAGE analysis, and endotoxin.

### Mouse Model of MODS

To model systemic inflammation associated with severe sepsis, nine-week-old wild-type female C57BL/6J mice (Jackson Laboratories) were immobilized via scruff before receiving an intraperitoneal (IP) injection of 10 mg/kg lipopolysaccharide (LPS) diluted at 1 mg/mL concentration in sterile saline. After 6 hours, mice received a secondary 1 mg/kg IP LPS injection. At 10 hours following the initial LPS injection, mice were arbitrarily selected for receiving either 200 μL of sterile saline or 10 mg/mL iECM injected into the tail vein with a 29G needle insulin syringe (Exel International, Redondo Beach, CA). Temperature and weight were taken preceding each handling timepoint. Any animals with a temperature above 28°C were excluded prior to injection with iECM or saline based on previous model validation showing this temperature profile indicated an inadequate inflammatory response to LPS. Treated animals were followed up for up to 20 hours post iECM or saline treatment to assess effect of the iECM material. All procedures in this study were approved by the Institutional Animal Care and Use Committee at the University of California San Diego, and performed according to guidelines from the Association for the Assessment and Accreditation of Laboratory Animal Care.

### Material Retention

To test the biodistribution of materials within the organs of LPS-induced mice, iECM or trilysine (Sigma-Aldrich) was tagged with FLUOR Vivotag-S 750 by NHS-ester chemistry as previously described (2) and delivered via 200 ul tail vein injection of a 10 mg/ml solution of iECM. Animals were arbitrarily selected and treated with either no LPS with iECM injection at 10 hours, LPS with iECM treatment, or LPS with trilysine at the same peptide concentration to iECM. Fluorescence intensity of multiple primary and satellite organs was measured using LiCor scanning after animal sacrifice at 14-hours. A total of n = 8-10 samples per organ per group were utilized. For each organ, the intensity normalized to area was calculated using MATLAB.

### Survival Analysis

All animals utilized for immunological characterization (ELISA, Nanostring, and flow cytometry) analyses were incorporated into the survival study for a total of n = 112 animals (n = 47 for iECM, n = 65 for saline). A higher n for saline was used to accommodate for the increased mortality. Survival curve analysis was performed using the Log-Rank (Mantel Cox) test.

### Multiplex ELISA

Animals treated with LPS and iECM or saline were sacrificed at hour 30 as previously described. A total of n = 6 animals were used per group for a total of 12 animals and whole blood, spleen, lung, heart, brain, and kidney were harvested and flash frozen. Protein lysate was isolated for each snap frozen tissue. Blood was collected in 1.5 mL K2 EDTA microtainer tubes (BD) and spun down at 1,400 RCF for 10 minutes to isolate plasma. Tissue samples were homogenized using a Beadbug bead homogenizer (Benchmark Scientific) in a complete extraction buffer composed of 100mM Tris (pH 7.4), 150 mM NaCl, 1 mM EGTA, 1 mM EDTA, 1% Triton X-100, 0.5% sodium deoxycholate, 1x phosSTOP (Sigma), 1x protease inhibitor cocktail (Sigma), and 1 mM PMSF with 60 μL of buffer used for every milligram of tissue. Homogenates were agitated at 4°C in a thermomixer (Eppendorf) for 2 hours, then centrifuged at 13,000 rpm at 4°C for 20 min. Supernatants were aliquoted on ice before storage at −80 °C. Supernatants were first used in a bicinchoninic acid (BCA) assay (Thermo Fisher) to identify the total protein in each extract. Supernatants were then assayed with a LEGENDPlex Mouse Inflammation Panel (13-plex, BioLegend) following the manufacturer’s protocol. Data was acquired using a BD LSRFortessaTM X-20 (BD Biosciences) followed by analysis in LEGENDplex V8.0 Data Analysis Software (BioLegend).

### RNA Isolation

Animals treated with LPS and iECM or saline were sacrificed at hour 30 as previously described. A total of n = 6 animals were used per group for a total of 12 animals and whole blood, spleen, lung, heart, brain, and kidney were harvested and flash frozen. 250 μL of whole blood was collected in 2 mL of RNAlater and stored at −80 °C until further processing. Flash frozen samples tissue samples were homogenized with either a Tissue Tearer rotator homogenizer (Cole-Parmer, Vernon Hills, IL) or a Beadbug bead homogenizer (Benchmark Scientific). For lung, brain, kidney, and spleen, RNA was isolated through a NucleoSpin® RNA/Protein kit (Takara Bio USA Inc. San Jose, CA) while heart issue was processed with a RNeasy Fibrous Mini kit (Qiagen, Germantown, MD). Blood samples were isolated using a Mouse Ribopure Blood RNA Isolation Kit (Invitrogen). RNA was extracted based on manufacturer instructions with an on-column DNase digestion for minimizing genomic DNA contamination.

### Nanostring nCounter Protocol

RNA samples were processed for Nanostring nCounter analysis according to manufacturer instructions. In brief, RNA sample concentrations were measured on a Qubit 3.0 Fluorometer with a QubitTM RNA HS Assay kit. 70 μL of hybridization buffer was mixed with Immunology Panel Reporter CodeSet solution, and 8 μL of this master mix was mixed in a separate reaction vessel with 100 ng of RNA per tissue sample and RNA-free water up to 13 μL total. 2 μL of Capture ProbeSet was added to each vessel, mixed and placed on a thermocycler at 65oC for 16-48 hours before being maintained at 4°C for less than 24 hours. Nanostring nCounter Prep Station performed automated fluidic sample processing to purify and immobilize hybridized sample to cartridge surface. Digital barcode reads were analyzed by Nanostring nCounter® Digital Analyzer.

### Nanostring Analysis via ROSALIND

Data was analyzed by ROSALIND®, with a HyperScale architecture developed by ROSALIND, Inc. (San Diego, CA). Read Distribution percentages, violin plots, identity heatmaps, and sample MDS plots were generated as part of the QC step. Normalization, fold changes and p-values were calculated using criteria provided by Nanostring. ROSALIND® follows the nCounter® Advanced Analysis protocol of dividing counts within a lane by the geometric mean of the normalizer probes from the same lane. Housekeeping probes to be used for normalization were selected based on the geNorm algorithm as implemented in the NormqPCR R library (3). Abundance of various cell populations was calculated on ROSALIND using the Nanostring Cell Type Profiling Module. In ROSALIND, we performed a filtering of Cell Type Profiling results to include results that have scores with a p-value greater than or equal to 0.05. Fold changes and p-values were calculated using the fast method as described in the nCounter® Advanced Analysis 2.0 User Manual. Pathway enrichment analysis was performed using the EnrichR package in R using the MSigDB Hallmark pathway library ^31^. Supplemental analysis was performed in PathfindR using the mouse WikiPathways pathway library^52^.

### Tissue Preparation for Spectral Flow Cytometry

The lungs of iECM or saline treated animals (n = 4 per treatment) were mechanically digested briefly prior to enzymatic digestion using a 0.5 mg/mL Liberase TM (Sigma) and a 0.2 mg/mL DNAse I (Sigma) digestion medium. Samples were enzymatically digested for 45 minutes at 37°C on a mechanical shaker before being strained with a 70 micron strainer and centrifuged. Samples were washed once in PBS, treated with 1x red blood cell lysis buffer (Biolegend), and washed two more times in 1x PBS. Samples were soaked in an EDTA solution for 10 minutes and resuspended in 1x PBS to a volume of 100 µl. Cells were stained with Live/DeadTM fixable blue (Invitrogen) to determine viability and then incubated with Monocyte blocking solution (Biolegend) to minimize off-target antibody binding. Cells were then washed in wash buffer (2mM EDTA and 1% BSA in 1x PBS) stained with a chemokine cocktail of anti-CX3CR1 and anti-XCR1 before addition of the main antibody cocktail containing the remainder of antibodies in **Supplemental Table 6**. Cells were washed and fixed using fixation buffer (Biolegend). Samples were kept overnight at 4°C and then run on a 5 Laser Cytek Aurora spectral cytometer. Cells were unmixed using the SpectroFlo software and then transferred to FlowJo for analysis.

### FlowJo Analysis of Spectral Flow Cytometry Samples

The gating strategy for flow cytometry is shown in **Supplementary Figure 15**. Samples were then concatenated into a single file and analyzed using flowSOM to identify unique populations^53^. The concatenated sample was initially visualized using Uniform Manifold Approximation and Projection (UMAP) for dimension reduction using the UMAP plugin in FlowJo. FlowSOM analysis was performed to identify 10 cell clusters present within all CD45+ cells in saline and iECM-treated animals. Heatmaps showing differential marker expression in clusters are visualized in **Supplementary Figure 17** and heatmaps of subclusters are in **Supplementary Figure 18.** Clusters are displayed in **Supplementary Figure 19** and subclusters are shown in **Supplementary Figure 20**. Samples were then analyzed through FlowSOM to visualize cell populations. Cell populations from gating in FlowJo were exported and transferred to Prism for visual representation and statistical analysis.

### Statistical Analysis

For two group assessments, an unpaired student’s t-test was performed while assessments across three or more groups was evaluated with a one-way ANOVA with a Tukey post-hoc with significance at p < 0.05. For analysis of differential expression of Nanostring data used a p value threshold of p < 0.05 and absolute fold change threshold of greater than or equal to 1.01. Gene set enrichment analysis utilized adjusted p values (padj) calculated using the Benjamini-Hochberg methodpadj < 0.05. Significant differences in survival curve were assessed using a Log-Rank (Mantel-Cox) test.

## Data Availability

Raw Nanostring files and analyses can be accessed on the GEO database (https://www.ncbi.nlm.nih.gov/geo/query/acc.cgi?acc=GSE268490, accession code sbgbwyaordofhed). Code for creation of volcano plots and pathway enrichment using PathfindR and EnrichR are detailed in Zenodo (https://zenodo.org/records/11290037?token=eyJhbGciOiJIUzUxMiJ9.eyJpZCI6ImM3YjBkNzcwLTE5Y2YtNDE4NC1hYjFkLWI4YmViZTkyYjlkYSIsImRhdGEiOnt9LCJyYW5kb20iOiJhZTYzZjY0YWYyNjVlZTJlOTFiODk5MTk1YzU2M2YyMiJ9.dOM3VJmnHm8gsl9rA61YAsmWpPBJFa0Xl7N9m4vACzi1lLEDxigw_ElW_g_DN6gOJ1lvY3x8t2w59x-V0FoqoQ).

## ACKNOWLEDGEMENTS

We would like to thank Dr. Elsa Molina at the UC San Diego Stem Cell Genomics Core, Stem Cell Program, La Jolla, California for her expertise and technical assistance with the NanoString nCounter assay. We would like to thank the LJI Flow Cytometry Core Facility for their technical expertise and assistance. Schematics in figures were created with BioRender. KLC is a co-founder, board member, and consultant and holds an equity interest in Ventrix Bio, Inc. All other authors have no relationships relevant to the contents of this paper. This work was supported by the California Institute for Regenerative Medicine (DISC2COVID19-12007 to KLC) and the National Institute of Health NHLBI (5R01HL165232 to KLC). Additional research support was provided by Department of Veterans Affairs IK2BX004338-01 (MLH) and the American Thoracic Society Research Foundation Unrestricted Critical Care Grant (MLH). AL and MK were supported by the National Science Foundation Graduate Research Fellowship (DGE-2038238 and 2022334614, respectively). J.M.M. and R.M.W. were supported by the National Institutes of Health (NIH), NHLBI Training Grant T32HL105373. A.C. was supported by the NIH, NIBIB Training Grant T32EB009380. JMM, AC, and MTS were supported by American Heart Association pre-doctoral fellowships.

