## Supplementary Materials for "An Infusible Extracellular Matrix Biomaterial Improves Survival in a Model of Severe Systemic Inflammation"

**Supplementary Table 1.** List of differentially expressed genes in whole blood of iECM treated animals versus saline.

| Symbol | log2FoldChange | pvalue |
|---|---|---|
| ABL1 | -1.87447 | 6.90E-04 |
| FOLR4 | -3.70044 | 7.22E-04 |
| CIITA | -2.78022 | 3.95E-05 |
| HAMP | -2.74982 | 3.66E-02 |
| MBL2 | -2.71786 | 7.78E-05 |
| TNFSF15 | -2.66297 | 1.29E-03 |
| TLR3 | -2.65992 | 3.65E-04 |
| MAPK11 | -2.65025 | 5.98E-04 |
| PSMB11 | -2.62449 | 2.00E-03 |
| RORC | -2.54889 | 4.21E-04 |
| CD59B | -2.54432 | 5.05E-03 |
| ADAL | -2.52083 | 2.06E-04 |
| CSF2 | -2.40599 | 2.52E-03 |
| MASP1 | -2.39855 | 1.07E-03 |
| NOX3 | -2.35279 | 9.13E-03 |
| CFI | -2.32193 | 3.30E-03 |
| TNFSF18 | -2.31159 | 8.24E-03 |
| IL2 | -2.30485 | 1.14E-02 |
| KIR3DL2 | -2.2878 | 1.37E-02 |
| NOX1 | -2.27798 | 2.82E-03 |
| MR1 | -2.27009 | 4.36E-03 |
| XCL1 | -2.19265 | 7.51E-03 |
| DEFB1 | -2.18133 | 7.44E-03 |
| CTSG | -2.17932 | 1.30E-02 |
| CCL11 | -2.17792 | 5.69E-03 |
| CR2 | -2.14505 | 1.14E-02 |
| CXCL15 | -2.14296 | 9.83E-03 |
| FCER1A | -2.1343 | 2.90E-03 |
| CXCR1 | -2.12553 | 1.55E-02 |
| RAG2 | -2.1247 | 1.02E-02 |
| PPARG | -2.12399 | 6.25E-03 |
| CD209G | -2.11548 | 8.80E-04 |
| GPR183 | -2.11306 | 1.21E-04 |
| KLRA6 | -2.10579 | 3.13E-03 |
| PLAU | -2.10236 | 5.67E-04 |
| IL23R | -2.09192 | 1.12E-02 |

| | | |
|---|---|---|
| GFI1 | -2.08746 | 4.83E-03 |
| PAX5 | -2.07973 | 8.16E-05 |
| KLRA1 | -2.07635 | 7.67E-03 |
| CLEC4A4 | -2.06301 | 1.34E-02 |
| IFNA2 | -2.05364 | 5.67E-03 |
| IL21 | -2.04041 | 2.06E-02 |
| AICDA | -2.02106 | 7.73E-03 |
| CD109 | -2 | 5.62E-03 |
| CD3EAP | -2 | 2.83E-03 |
| NFATC2 | -1.97461 | 5.92E-04 |
| SRC | -1.97199 | 1.75E-02 |
| CD5 | -1.96875 | 8.15E-04 |
| PDGFRB | -1.94251 | 5.03E-03 |
| IKZF3 | -1.92246 | 1.49E-05 |
| TNFRSF4 | -1.89924 | 5.95E-03 |
| CCL25 | -1.88501 | 1.19E-02 |
| IRF3 | -1.86836 | 1.23E-02 |
| NCAM1 | -1.86507 | 9.97E-03 |
| BATF3 | -1.8625 | 6.16E-03 |
| IL33 | -1.8609 | 4.86E-03 |
| CX3CR1 | -1.8606 | 1.15E-02 |
| STAT4 | -1.85914 | 1.81E-04 |
| IL27 | -1.85561 | 1.17E-02 |
| KLRA7 | -1.85561 | 7.99E-04 |
| PDGFB | -1.85561 | 2.55E-02 |
| CCR6 | -1.85244 | 1.39E-03 |
| CD8B1 | -1.83458 | 1.91E-02 |
| IL18 | -1.81214 | 3.48E-04 |
| FOXP3 | -1.8011 | 2.82E-02 |
| IL19 | -1.79518 | 3.50E-02 |
| IL23A | -1.78932 | 8.96E-03 |
| CX3CL1 | -1.78427 | 1.30E-02 |
| PLA2G2A | -1.75845 | 2.41E-02 |
| PDCD1 | -1.75207 | 4.22E-03 |
| FADD | -1.74543 | 3.75E-03 |
| IL7 | -1.74543 | 5.44E-03 |
| PDCD1LG2 | -1.72583 | 1.11E-02 |
| CD1D1 | -1.72403 | 1.37E-03 |
| TSLP | -1.71621 | 1.12E-02 |
| CXCL11 | -1.70044 | 5.48E-03 |

| | | |
|---|---|---|
| IL27RA | -1.68706 | 1.60E-02 |
| TNFSF11 | -1.6845 | 1.64E-02 |
| HFE | -1.654 | 4.20E-02 |
| CD8A | -1.64211 | 1.48E-02 |
| CD27 | -1.64111 | 2.97E-02 |
| PIGR | -1.63691 | 1.60E-02 |
| CCR3 | -1.6172 | 2.73E-02 |
| TBX21 | -1.61667 | 1.65E-02 |
| ZEB1 | -1.61257 | 8.43E-03 |
| CD3E | -1.60486 | 2.86E-02 |
| TNFRSF11A | -1.60192 | 2.77E-02 |
| CD163 | -1.58496 | 1.53E-02 |
| CXCR3 | -1.58496 | 1.55E-02 |
| IL12RB1 | -1.58496 | 1.81E-02 |
| TLR1 | -1.58496 | 1.06E-02 |
| DPP4 | -1.57491 | 2.32E-03 |
| IL9 | -1.56457 | 4.17E-02 |
| LILRA6 | -1.5502 | 3.60E-02 |
| LILRA5 | -1.54543 | 1.47E-02 |
| LIF | -1.54124 | 2.51E-02 |
| TNFRSF17 | -1.54057 | 4.23E-02 |
| IL12B | -1.53797 | 2.22E-02 |
| IKZF2 | -1.53303 | 3.66E-02 |
| KLRD1 | -1.52356 | 3.70E-02 |
| CD53 | -1.51679 | 1.68E-03 |
| PHLPP2 | -1.51525 | 2.31E-03 |
| XCR1 | -1.50696 | 1.20E-02 |
| IL25 | -1.50611 | 2.05E-02 |
| ILF3 | -1.46949 | 5.11E-03 |
| MX1 | -1.46546 | 2.19E-02 |
| C6 | -1.44815 | 3.45E-02 |
| STAT2 | -1.4458 | 2.17E-02 |
| TYK2 | -1.44057 | 2.20E-03 |
| C8B | -1.42321 | 2.33E-02 |
| CCL8 | -1.42084 | 1.92E-02 |
| TRAF1 | -1.41504 | 2.05E-02 |
| GATA3 | -1.38702 | 4.36E-02 |
| CCL24 | -1.37851 | 2.87E-02 |
| TRAF3 | -1.37266 | 3.91E-03 |
| IKBKB | -1.35216 | 2.46E-05 |

| | | |
|---|---|---|
| C9 | -1.34792 | 4.99E-02 |
| EOMES | -1.32193 | 2.05E-02 |
| TREM2 | -1.32193 | 2.63E-02 |
| GPI1 | -1.30319 | 1.92E-02 |
| LCP2 | -1.29336 | 1.30E-02 |
| KLRA8 | -1.27462 | 4.57E-02 |
| RUNX3 | -1.27085 | 4.33E-02 |
| PHLPP1 | -1.2655 | 1.31E-03 |
| CSF1 | -1.2645 | 1.49E-02 |
| GM10499 | -1.24236 | 3.32E-03 |
| CCL19 | -1.24042 | 1.57E-02 |
| IL17F | -1.18903 | 3.48E-02 |
| SMAD5 | -1.16651 | 4.37E-02 |
| C2 | -1.14886 | 2.15E-02 |
| PRDM1 | -1.13588 | 4.24E-02 |
| STAT5A | -1.13358 | 2.00E-02 |
| MASP2 | -1.10434 | 3.74E-02 |
| IL1R1 | -1.09954 | 1.62E-02 |
| CRADD | -1.0915 | 2.25E-02 |
| CASP2 | -1.08607 | 3.85E-02 |
| HCST | -1.08342 | 2.01E-02 |
| TNFRSF13C | -1.02647 | 3.05E-02 |
| TRAF5 | -1.02086 | 6.59E-03 |
| IL18R1 | -0.99512 | 2.91E-02 |
| CCR9 | -0.96919 | 1.09E-02 |
| CD86 | -0.93449 | 1.43E-02 |
| MS4A1 | -0.9 | 4.19E-02 |
| CFH | -0.86135 | 4.07E-02 |
| NOTCH1 | -0.8593 | 1.67E-02 |
| TRAF6 | -0.85775 | 3.09E-02 |
| IL1A | -0.85609 | 3.44E-02 |
| TAGAP | -0.8501 | 3.03E-02 |
| TGFBR1 | -0.81431 | 4.89E-02 |
| CD80 | -0.80512 | 6.58E-03 |
| IL6 | -0.79105 | 2.89E-03 |
| IKZF1 | -0.77091 | 1.29E-02 |
| IL15RA | -0.72289 | 3.73E-02 |
| TCF4 | -0.69341 | 4.13E-02 |
| ITGA4 | -0.69285 | 2.43E-02 |
| BTK | -0.66896 | 3.91E-02 |

| | | |
|---|---|---|
| SKI | -0.66051 | 2.56E-02 |
| RAE1 | -0.62207 | 2.23E-02 |
| IFNGR2 | -0.60768 | 4.95E-02 |
| NPC1 | -0.58496 | 4.63E-03 |
| ATG16L1 | -0.53222 | 3.29E-02 |
| ICAM1 | -0.48917 | 3.92E-03 |
| NFIL3 | -0.47286 | 4.60E-02 |
| CASP8 | -0.3647 | 2.79E-02 |
| TAPBP | 0.335239 | 2.47E-02 |
| CD164 | 0.410376 | 3.45E-02 |
| CRLF2 | 0.42752 | 1.82E-02 |
| IL10RB | 0.467505 | 4.01E-02 |
| STAT1 | 0.483724 | 3.18E-03 |
| CD97 | 0.485427 | 3.83E-02 |
| TNFRSF1B | 0.494872 | 3.19E-02 |
| BCAP31 | 0.523445 | 2.70E-02 |
| SELL | 0.545412 | 2.01E-02 |
| IFNAR2 | 0.644067 | 2.09E-02 |
| IFI35 | 0.693215 | 1.11E-02 |
| FCGR4 | 0.697636 | 1.18E-02 |
| CASP3 | 0.736019 | 4.55E-03 |
| FCGR3 | 0.772674 | 3.48E-02 |
| IFNGR1 | 0.824104 | 4.46E-03 |
| B2M | 0.833421 | 1.58E-04 |
| EBI3 | 0.876039 | 5.89E-03 |
| IFI204 | 0.878371 | 6.18E-03 |
| TGFB1 | 0.900395 | 7.27E-03 |
| TYROBP | 0.942191 | 1.62E-02 |
| CD82 | 0.961497 | 3.15E-02 |
| IFITM1 | 0.991222 | 3.26E-02 |
| PTPN6 | 1.014229 | 3.90E-03 |
| CD81 | 1.063706 | 6.75E-04 |
| ARHGDIB | 1.102635 | 1.01E-03 |
| NCF4 | 1.145198 | 1.30E-03 |
| IL1R2 | 1.171469 | 4.21E-02 |
| KLRK1 | 1.344383 | 1.46E-02 |
| S100A9 | 1.424649 | 4.80E-02 |
| PDCD2 | 1.448758 | 3.63E-02 |
| S100A8 | 1.488409 | 3.25E-02 |
| CD9 | 1.516765 | 2.67E-02 |

| | | |
|---|---|---|
| HLX | 1.603341 | 5.84E-03 |
| ITGA2B | 1.615551 | 4.50E-02 |
| PPBP | 1.785991 | 3.35E-02 |
| GZMA | 2.125531 | 4.18E-05 |
| CARD9 | 2.688056 | 3.49E-04 |

**Supplementary Table 2**. List of differentially expressed genes in brain tissue of iECM treated animals versus saline.

| Symbol | log2FoldChange | pvalue |
|---|---|---|
| **IL6** | -1.4394 | 0.000298 |
| DEFB1 | -1.37587 | 0.014572 |
| IL19 | -1.01992 | 0.004209 |
| CXCL11 | -1.01668 | 0.014009 |
| CCL2 | -0.96722 | 0.020033 |
| CXCL1 | -0.91519 | 0.004728 |
| CCL7 | -0.89812 | 0.024907 |
| H2-AB1 | -0.89121 | 0.003442 |
| H2-AA | -0.82238 | 0.0099 |
| CCL11 | -0.70751 | 0.045507 |
| LCK | -0.70711 | 0.020802 |
| IL1RN | -0.69401 | 0.039864 |
| CCL5 | -0.64088 | 0.004004 |
| CD74 | -0.61268 | 0.000433 |
| CD80 | -0.57308 | 0.004023 |
| CCL9 | -0.52575 | 0.013073 |
| RUNX1 | -0.51802 | 0.005683 |
| VCAM1 | -0.47366 | 0.016621 |
| IL15RA | -0.47256 | 0.006992 |
| ICAM1 | -0.45701 | 0.015312 |
| CD274 | -0.45298 | 0.019272 |
| IFIT2 | -0.437 | 0.026013 |
| NT5E | -0.41734 | 0.028939 |
| IFITM1 | -0.39089 | 0.039581 |
| CD109 | -0.38464 | 0.030325 |
| TLR2 | -0.37599 | 0.038863 |
| NFKB2 | -0.3619 | 0.006074 |
| IL13RA1 | -0.35571 | 0.007082 |
| IFIH1 | -0.35551 | 0.021024 |
| TAP1 | -0.35528 | 0.002343 |
| CASP8 | -0.31989 | 0.04667 |
| SOCS3 | -0.31905 | 0.041821 |
| PLAUR | -0.31715 | 0.024903 |
| FAS | -0.31352 | 0.031958 |
| NOTCH2 | -0.29189 | 0.039626 |
| ETS1 | -0.29125 | 0.004875 |
| IRAK3 | -0.24246 | 0.034115 |

| | | |
|---|---|---|
| CCL25 | -0.22529 | 0.009238 |
| NCAM1 | -0.17067 | 0.045459 |
| PECAM1 | 0.282955 | 0.029562 |
| TFRC | 0.3181 | 0.003679 |
| MAPK11 | 0.386417 | 0.006069 |
| STAT6 | 0.430325 | 0.033148 |
| LAIR1 | 0.515133 | 0.003405 |
| TRAF5 | 0.516454 | 0.01793 |
| IL1RL2 | 0.555816 | 0.022307 |
| ICAM2 | 0.571906 | 0.034015 |
| CD48 | 1.039966 | 0.007231 |
| IL12A | 1.106915 | 0.024603 |
| CX3CR1 | 1.117326 | 0.037299 |
| CD34 | 1.303071 | 0.007096 |
| PPARG | 1.329214 | 0.002961 |

**Supplementary Table 3.** List of differentially expressed genes in lung tissue of iECM treated animals versus saline.

| Symbol | log2FoldChange | pvalue |
|---|---|---|
| **IL17A** | -1.66558 | 0.03973 |
| LIF | -1.3694 | 0.01396 |
| IL6 | -1.26052 | 0.021344 |
| CXCL3 | -1.22257 | 0.00178 |
| TSLP | -1.05557 | 0.000307 |
| IL19 | -1.03747 | 0.039501 |
| CXCL1 | -0.99573 | 0.033263 |
| TNFRSF8 | -0.91754 | 0.005975 |
| TNFSF15 | -0.89308 | 0.023618 |
| IL15 | -0.86953 | 0.042718 |
| CCL4 | -0.85293 | 0.00084 |
| PTGS2 | -0.82525 | 0.012407 |
| IL1A | -0.82433 | 0.02464 |
| TNFAIP6 | -0.80084 | 0.02287 |
| CTLA4 | -0.76268 | 0.013044 |
| CD14 | -0.70748 | 0.000372 |
| NFIL3 | -0.70364 | 0.010672 |
| IL12RB2 | -0.68575 | 0.002164 |
| IRF4 | -0.6734 | 0.03384 |
| TNFRSF9 | -0.64261 | 0.005565 |
| CCR9 | -0.63912 | 0.025027 |
| CXCL15 | -0.62679 | 0.00093 |
| HIF1A | -0.62286 | 0.000529 |
| VCAM1 | -0.60987 | 0.018383 |
| SOCS3 | -0.58862 | 0.04346 |
| MX1 | -0.58758 | 0.015243 |
| PML | -0.58128 | 0.00782 |
| CISH | -0.56954 | 0.001114 |
| PRDM1 | -0.5121 | 0.006677 |
| LITAF | -0.51057 | 0.001973 |
| ETS1 | -0.49198 | 0.004077 |
| CCL3 | -0.47459 | 0.040118 |
| CD53 | -0.46287 | 0.041742 |
| ENTPD1 | -0.43954 | 0.010647 |
| PTPN2 | -0.41663 | 0.026334 |
| IRF8 | -0.41378 | 0.020894 |
| CCL5 | -0.4026 | 0.019108 |

| | | |
|---|---|---|
| TLR8 | -0.38428 | 0.019829 |
| CEBPB | -0.36376 | 0.013002 |
| BCL6 | -0.34776 | 0.004155 |
| CTSC | -0.33497 | 0.002236 |
| CD44 | -0.33295 | 0.017753 |
| MAPKAPK2 | -0.31948 | 0.001274 |
| STAT3 | -0.30506 | 0.001545 |
| CCL9 | -0.29374 | 0.034475 |
| GM10499 | -0.29273 | 0.009283 |
| MYD88 | -0.29002 | 0.004405 |
| RUNX1 | -0.27022 | 0.037664 |
| JAK1 | -0.24505 | 0.010406 |
| TFRC | -0.23494 | 0.014161 |
| IFNAR2 | -0.23217 | 0.040449 |
| PSMB7 | -0.23158 | 0.01496 |
| NFKB1 | -0.22703 | 0.034519 |
| TGFB1 | -0.22152 | 0.029722 |
| ABCF1 | -0.20812 | 0.020808 |
| TGFBR1 | -0.15646 | 0.034663 |
| IRAK1 | 0.182711 | 0.047391 |
| FCGRT | 0.320237 | 0.013847 |
| SIGIRR | 0.371811 | 0.022527 |
| PDGFB | 0.386251 | 0.038447 |
| IL17RE | 0.40691 | 0.046482 |
| MR1 | 0.563901 | 0.024897 |
| FCAMR | 0.564953 | 0.020124 |
| CXCL13 | 0.572713 | 0.013897 |
| MAP4K2 | 0.589354 | 0.006868 |
| CMKLR1 | 0.623237 | 0.001189 |
| H2-OB | 0.736966 | 0.031374 |
| CD163 | 0.858927 | 0.002812 |
| IL11RA1 | 0.928107 | 0.027597 |

**Supplementary Table 4.** List of differentially expressed genes in heart tissue of iECM treated animals versus saline.

| Symbol | log2FoldChange | pvalue |
|---|---|---|
| PTGS2 | -1.72059 | 0.005579 |
| IL6 | -1.3833 | 0.014042 |
| CD24A | -0.93032 | 0.012996 |
| CXCL1 | -0.90129 | 0.018418 |
| TNFAIP6 | -0.86964 | 0.014337 |
| SOCS3 | -0.70344 | 0.032569 |
| CD14 | -0.60231 | 0.000378 |
| PLAUR | -0.48501 | 0.006499 |
| NFKBIZ | -0.43522 | 0.010382 |
| LITAF | -0.30337 | 0.021334 |
| CEBPB | -0.28226 | 0.011443 |
| CDH5 | -0.22198 | 0.02238 |
| IRAK3 | -0.21229 | 0.034542 |
| PHLPP2 | -0.1735 | 0.047316 |
| PTPN2 | -0.154 | 0.003312 |
| PPARG | 0.248873 | 0.03739 |
| ABCB1A | 0.259558 | 0.027228 |
| PTPN6 | 0.313557 | 0.03019 |
| CD3EAP | 0.356144 | 0.037029 |

**Supplementary Table 5.** List of differentially expressed genes in kidney tissue of iECM treated animals versus saline.

| Symbol | log2FoldChange | pvalue |
|---|---|---|
| C6 | 0.625073 | 0.029272 |
| ICOS | -1.55459 | 0.001547 |
| CCL20 | -1.01073 | 0.046332 |
| IL1B | -0.82912 | 0.011872 |
| TNFAIP6 | -0.81073 | 0.027712 |
| CXCL11 | -0.80979 | 0.027433 |
| CD80 | -0.65208 | 0.039079 |
| CXCL1 | -0.61953 | 0.033337 |
| CD44 | -0.56084 | 0.016704 |
| IL1RN | -0.416 | 0.047098 |
| H2-AB1 | 0.245338 | 0.042242 |
| CD34 | 0.257323 | 0.026991 |
| PTPN6 | 0.287697 | 0.04075 |
| TRAF5 | 0.484428 | 0.002403 |
| ITGB2 | 0.643451 | 0.015688 |
| LY86 | 0.68281 | 0.038017 |
| C7 | 0.795662 | 0.000992 |
| CD247 | 1.242857 | 0.003677 |
| CD79B | 1.242857 | 0.034772 |
| CCL8 | 1.484561 | 0.013577 |

**Supplementary Table 6.** Overview of flow cytometry antibodies used for a 22 color myeloid cell profile.

| Fluorophore | Target | Company |
|---|---|---|
| BUV395 | CD45 | BD |
| LIVE DEAD Fixable Blue | Dead Cells | Thermo |
| BUV 496 | MHCII | BD |
| BUV563 | CD1d | BD |
| BUV805 | F4/80 | BD |
| BV421 | CD206 | Biolegend |
| eFluor 450 | Ly6G | Thermo |
| BV480 | CD103 | BD |
| BV510 | Ly6C | Biolegend |
| BV570 | CD11c | Biolegend |
| BV605 | Siglec-F | BD |
| BV650 | XCR1 | Biolegend |
| FITC | CD43 | Biolegend |
| Spark Blue 550 | CD8α | Biolegend |
| PE/Cy7 | CD301b | Biolegend |
| PE | CD86 | Biolegend |
| PE/Dazzle594 | CD169 | Biolegend |
| PE-Cy5 | CCR7 | Biolegend |
| APC | CD200R3 | Biolegend |
| Alexa Fluor 647 | CX3CR1 | Biolegend |
| Alexa Fluor 700 | CD11b | Biolegend |
| APC-Cy7 | CD68 | Biolegend |

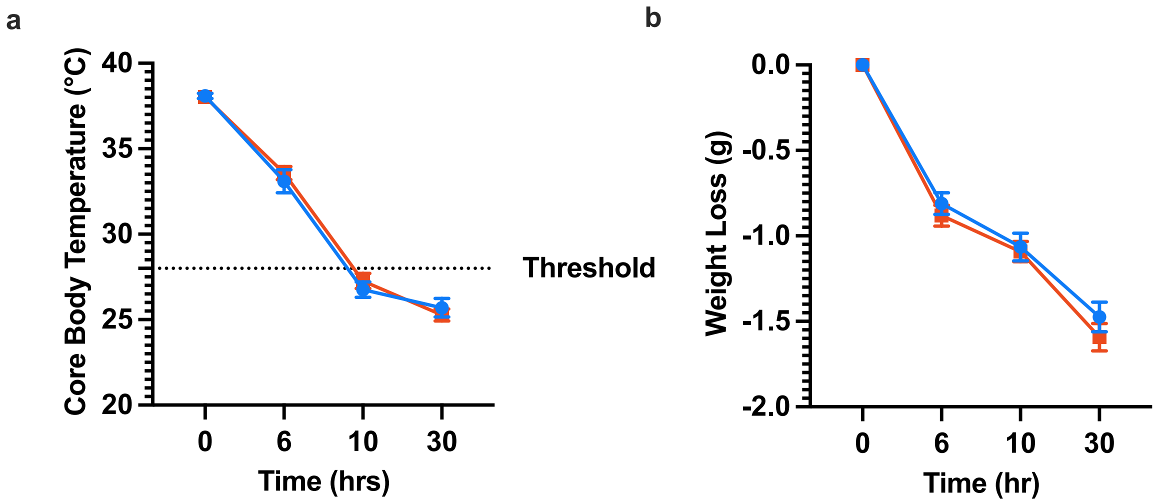


**Supplemental Figure 1. Dynamics of (a) core body temperature and (b) weight drop in MODS animals treated with iECM (blue) or saline (red) over a 30 hour period**. Dotted line in (**a**) represents the threshold temperature at 10 hours below that determined successful onset of MODS and dysregulation of homeostasis.

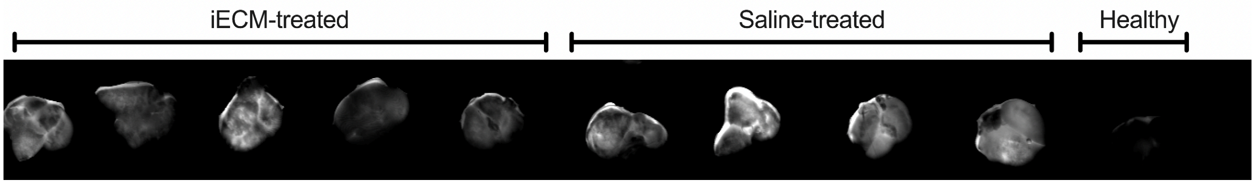


**Supplemental Figure 2. Raw image of lung tissue showing retention of fluorescently labelled bovine-serum albumin (BSA).**

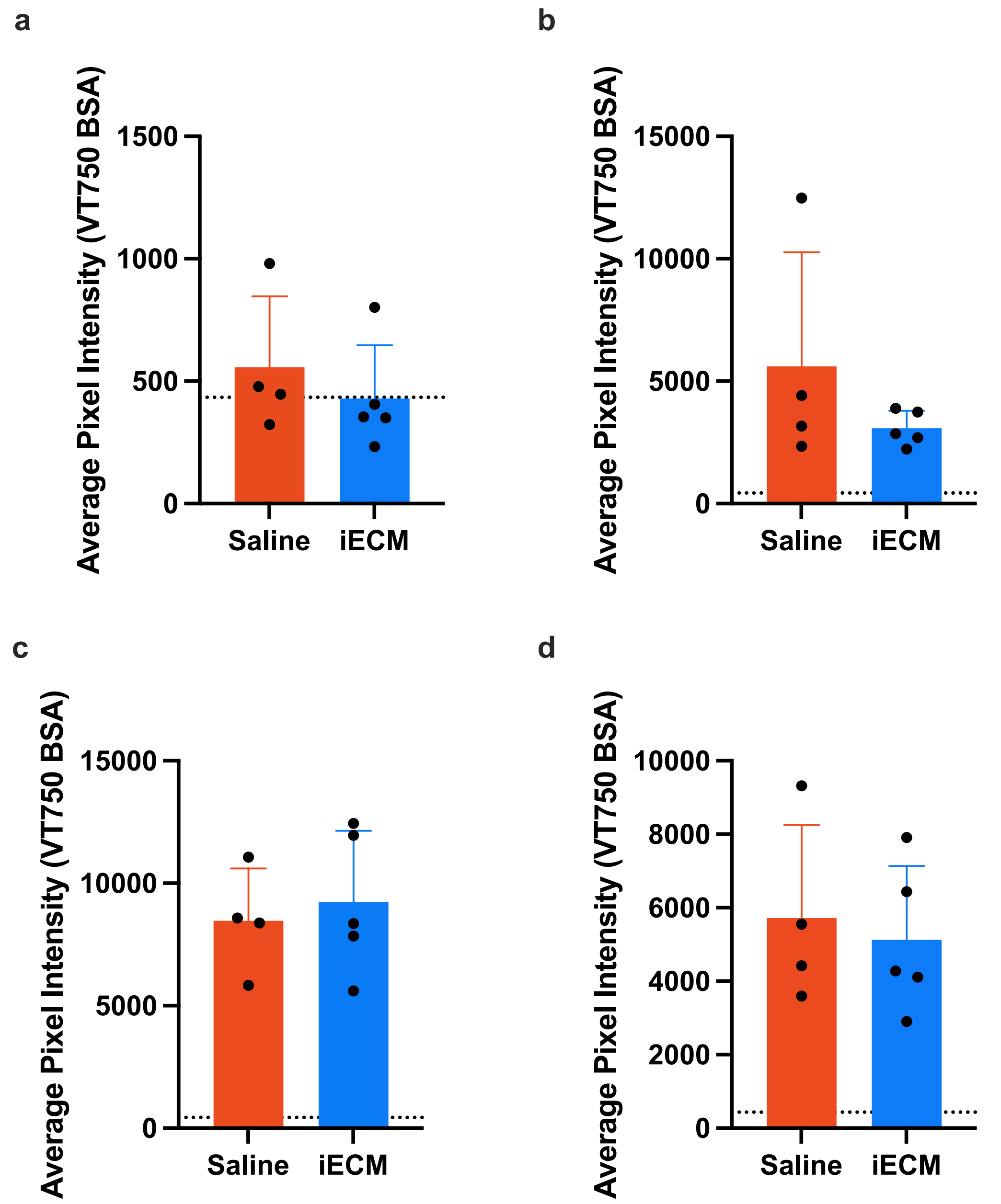


**Supplemental Figure 3. Quantification of fluorescently-labelled albumin (VT750 BSA) in (a) brain (b) heart (c) kidney and (d) liver of animals treated with saline (red) or iECM (blue).** Animals were treated with two doses of LPS, and then treated with iECM or saline 10 hours after initial LPS dose. One hour after iECM or saline administration, fluorescently labelled albumin was injected intravenously. Animals were perfused with 1x PBS and harvested 1 hour post albumin injection; organs were imaged on a Licor Odyssey to visualize albumin leakage in tissues. Average pixel intensity was determined in FIJI and significance was determined using a two-tailed unpaired t-test. P* <0.05. Dotted line represents baseline levels of fluorescent signal in healthy animals injected with fluorescently-labelled albumin.

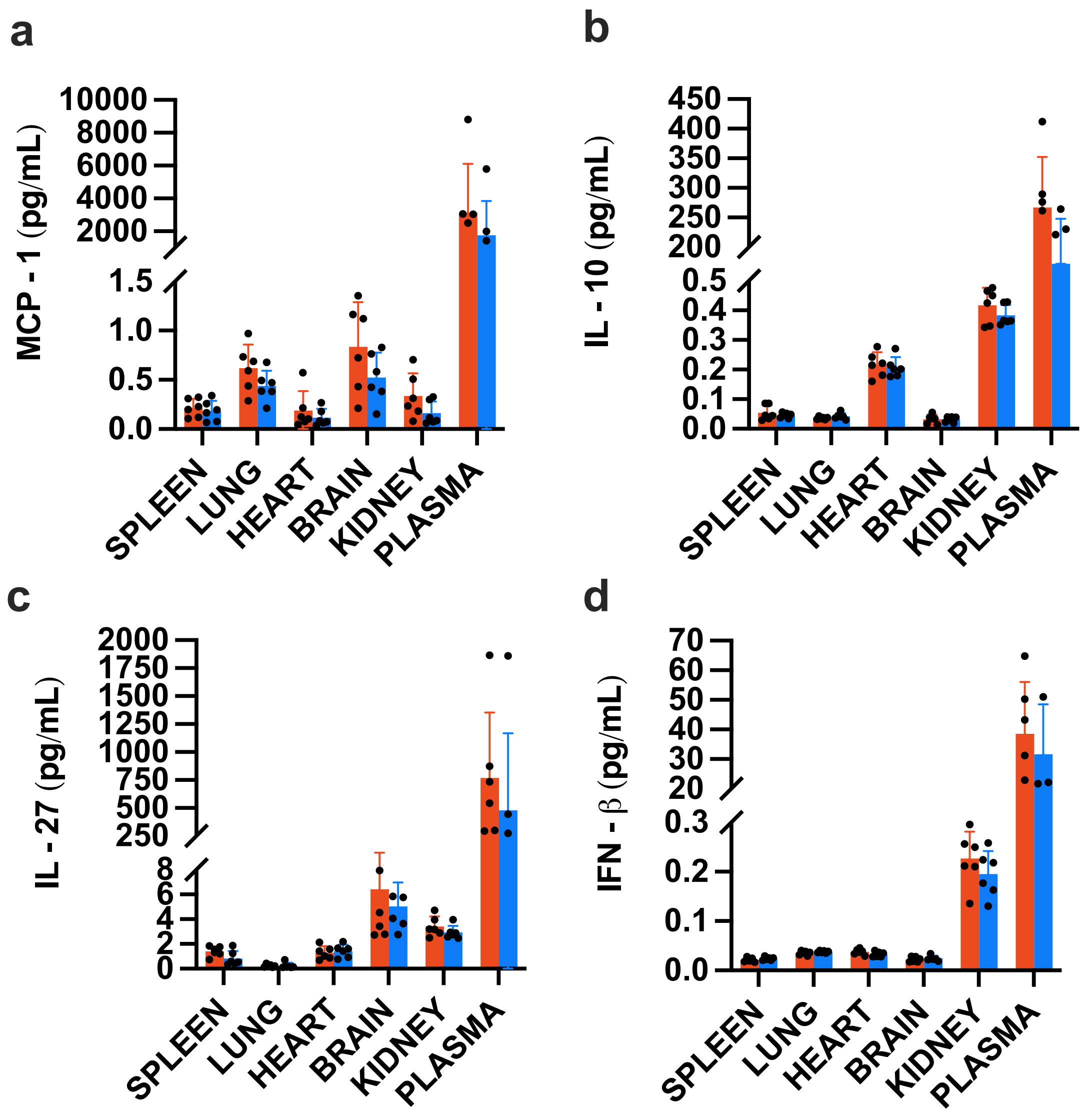


**Supplemental Figure 4. Levels of (a) MCP-1, (b) IL-10, (c) IL-27, and (d) IFN-β probed in saline (red) and iECM (blue)-treated animals (n = 6 per group) using a 13-plex ELISA array (Biolegend).** Significance was determined using an unpaired t-test with individual variance assumption for each row to accommodate for inherent differences in baseline tissue expression of each cytokine.

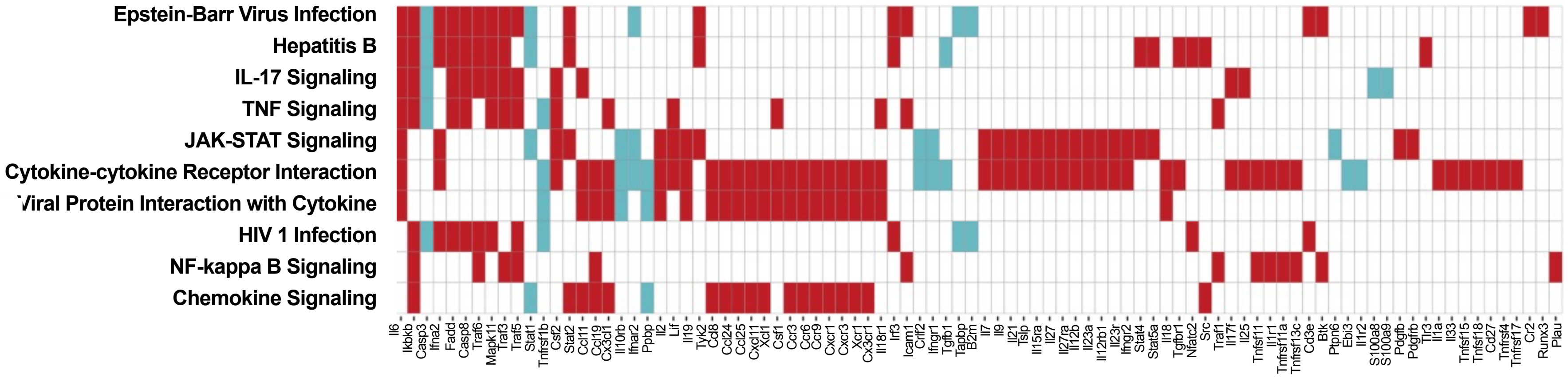


**Supplementary Figure 5. Heatmap of pathway enrichment analysis using PathfindR for differentially expressed genes (upregulated = teal, downregulated = red) in blood.** A total of n = 5-6 RNA samples per tissue were run on the Nanostring nCounter Immunology panel. Differential expression was determined using ROSALIND. Differentially expressed genes were inputted into R and analyzed using pathfindR.

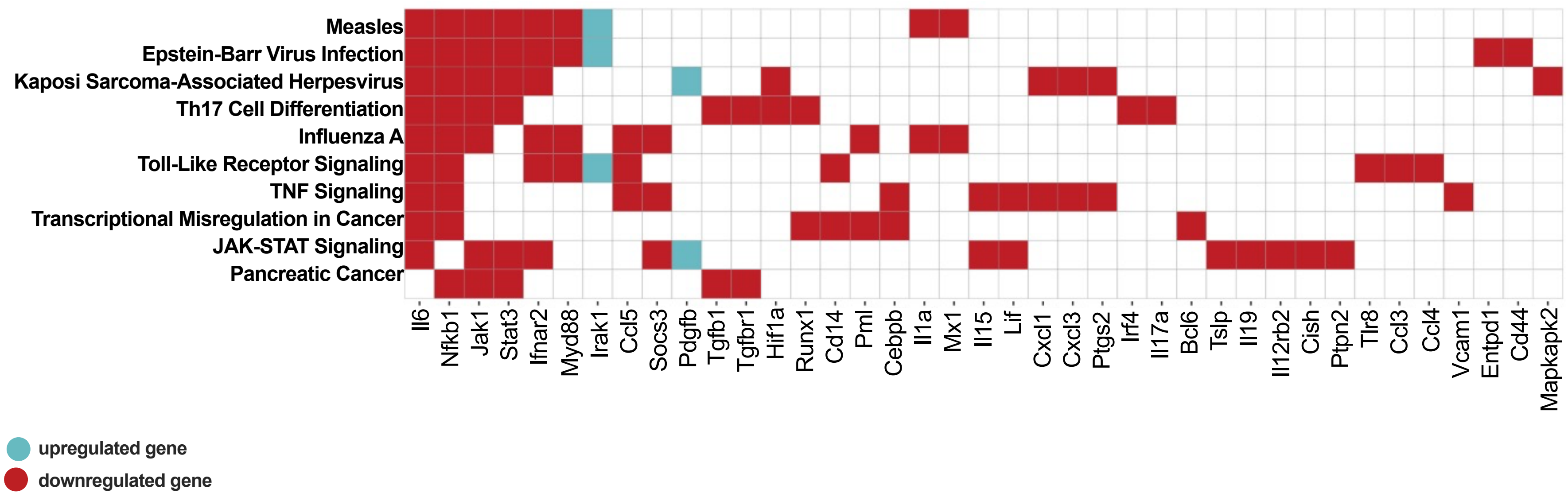


**Supplementary Figure 5. Heatmap of pathway enrichment analysis using PathfindR for differentially expressed genes (upregulated = teal, downregulated = red) in lung tissue.** A total of n = 5-6 RNA samples per tissue were run on the Nanostring nCounter Immunology panel. Differential expression was determined using ROSALIND. Differentially expressed genes were inputted into R and analyzed using pathfindR.

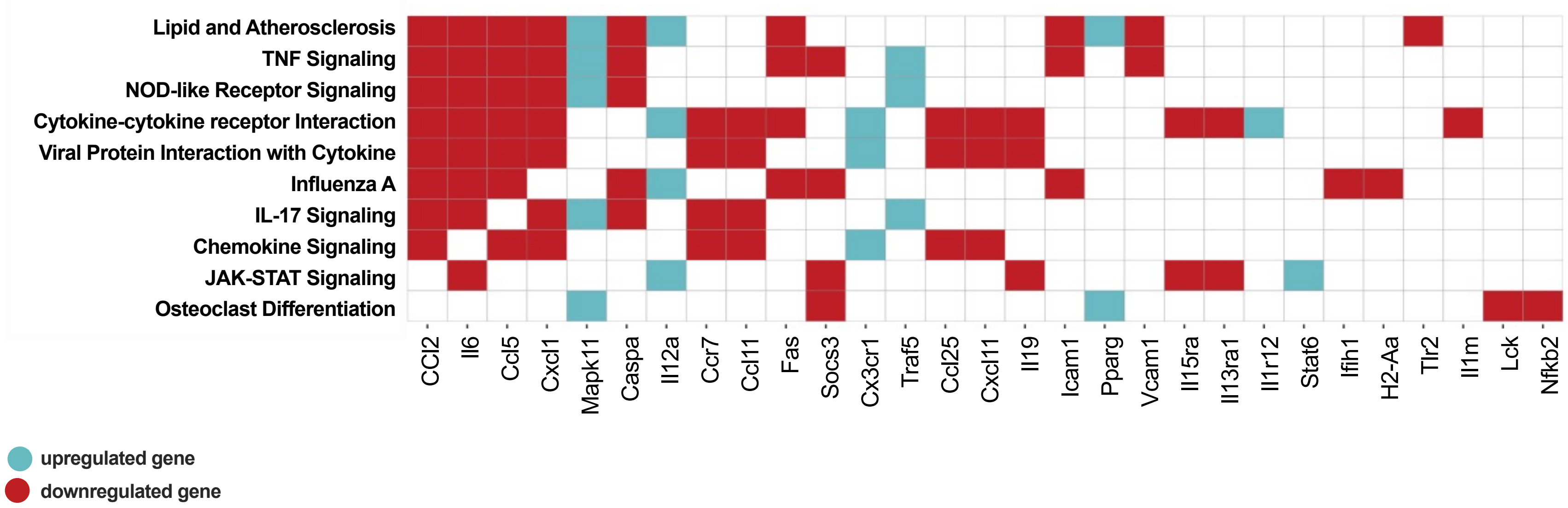


**Supplemental Figure 7. Heatmap of pathway enrichment analysis using PathfindR for differentially expressed genes (upregulated = teal, downregulated = red) in brain tissue.** A total of n = 5-6 RNA samples per tissue were run on the Nanostring nCounter Immunology panel. Differential expression was determined using ROSALIND. Differentially expressed genes were inputted into R and analyzed using pathfindR.

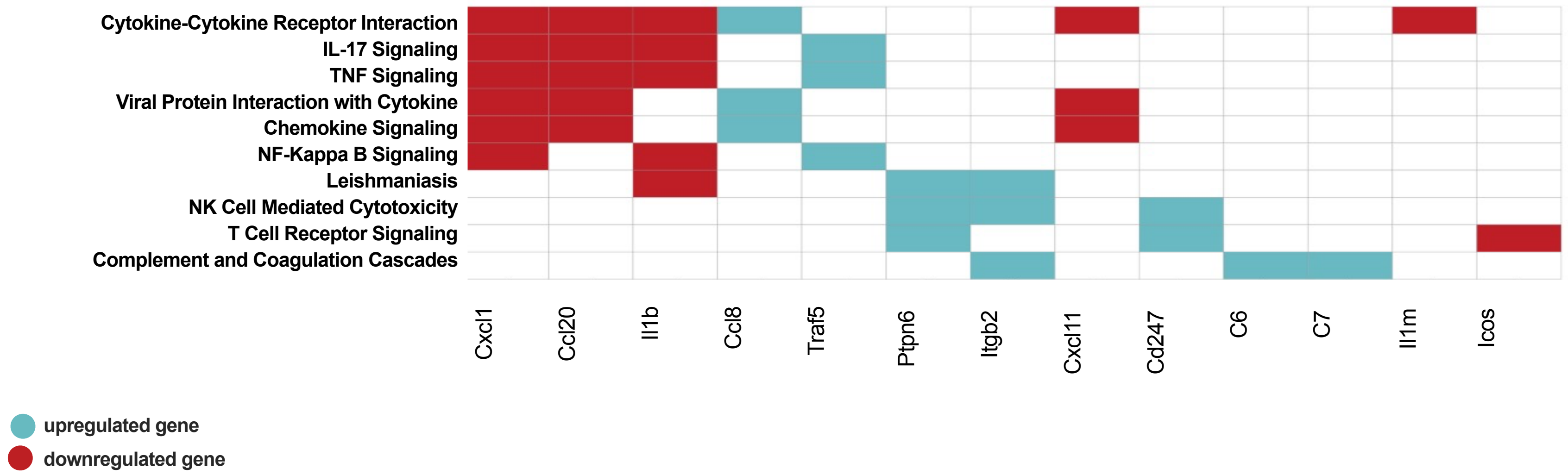


**Supplemental Figure 8. Heatmap of pathway enrichment analysis using PathfindR for differentially expressed genes (upregulated = teal, downregulated = red) in kidney tissues.** A total of n = 5-6 RNA samples per tissue were run on the Nanostring nCounter Immunology panel. Differential expression was determined using ROSALIND. Differentially expressed genes were inputted into R and analyzed using pathfindR.

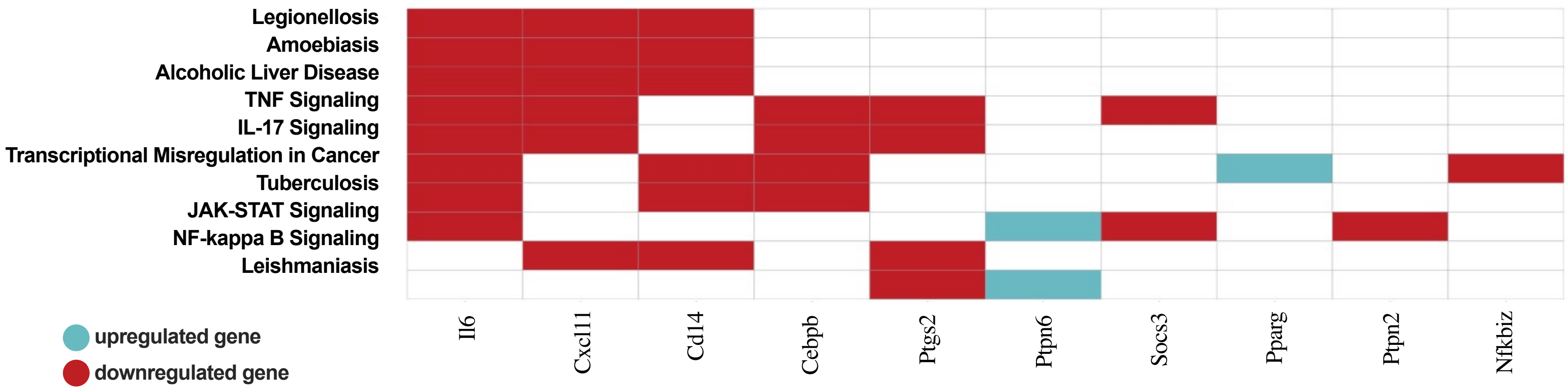


**Supplemental Figure 9. Heatmap of pathway enrichment analysis using PathfindR for differentially expressed genes (upregulated = teal, downregulated = red) in heart tissue.** A total of n = 5-6 RNA samples per tissue were run on the Nanostring nCounter Immunology panel. Differential expression was determined using ROSALIND. Differentially expressed genes were inputted into R and analyzed using pathfindR.

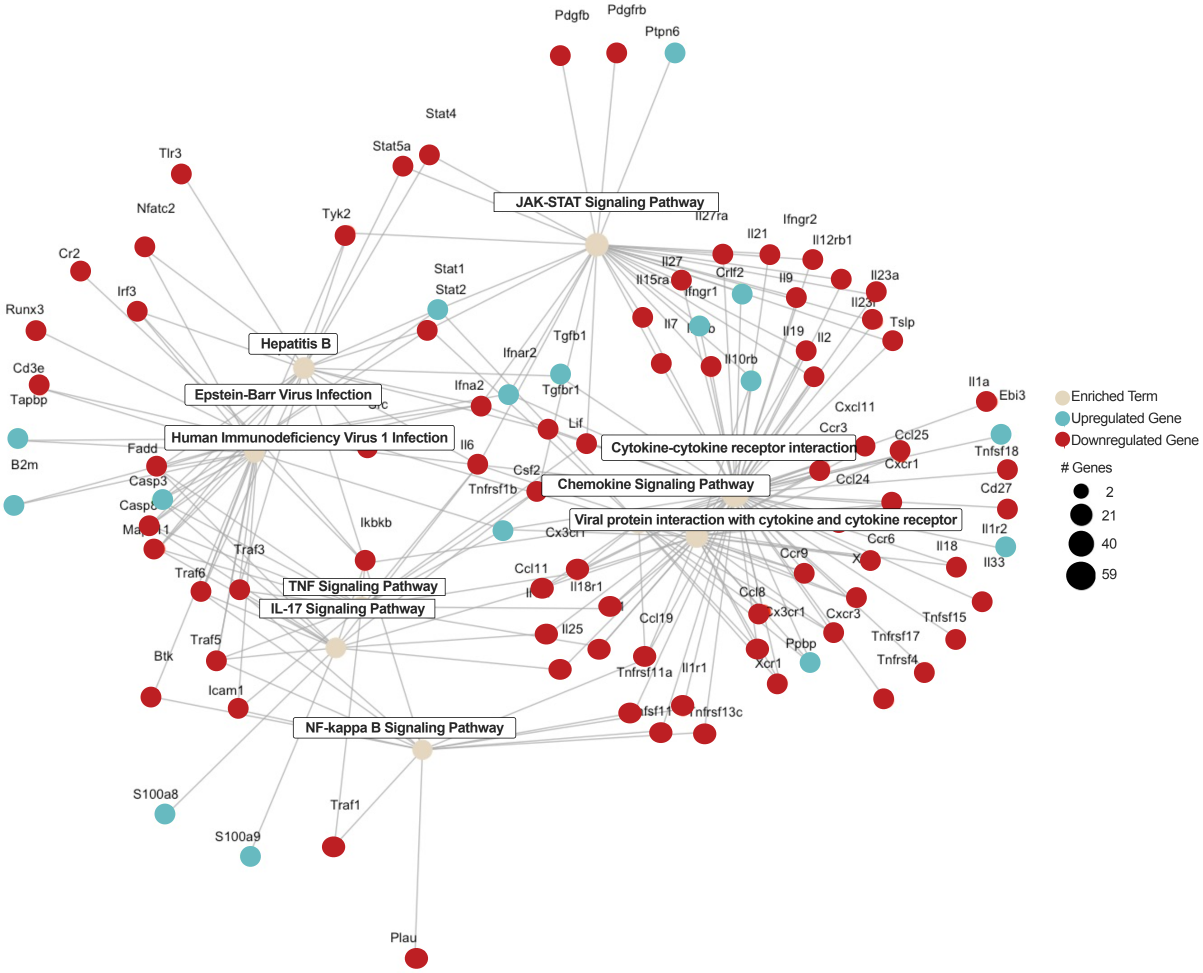


**Supplemental Figure 10. Gene map of significantly differentially expressed genes ($p < 0.05$) in blood samples in iECM vs saline-treated animals.** A total of n = 5 blood RNA samples were utilized and run on the Nanostring nCounter panel. Differential expression was determined using ROSALIND. Differentially expressed genes were inputted into R and analyzed using pathfindR. Size of enriched term node is correlated to number of differentially expressed genes within that pathway. Genes upregulated with iECM treatment are shown in blue, while downregulated genes are visualized in red.

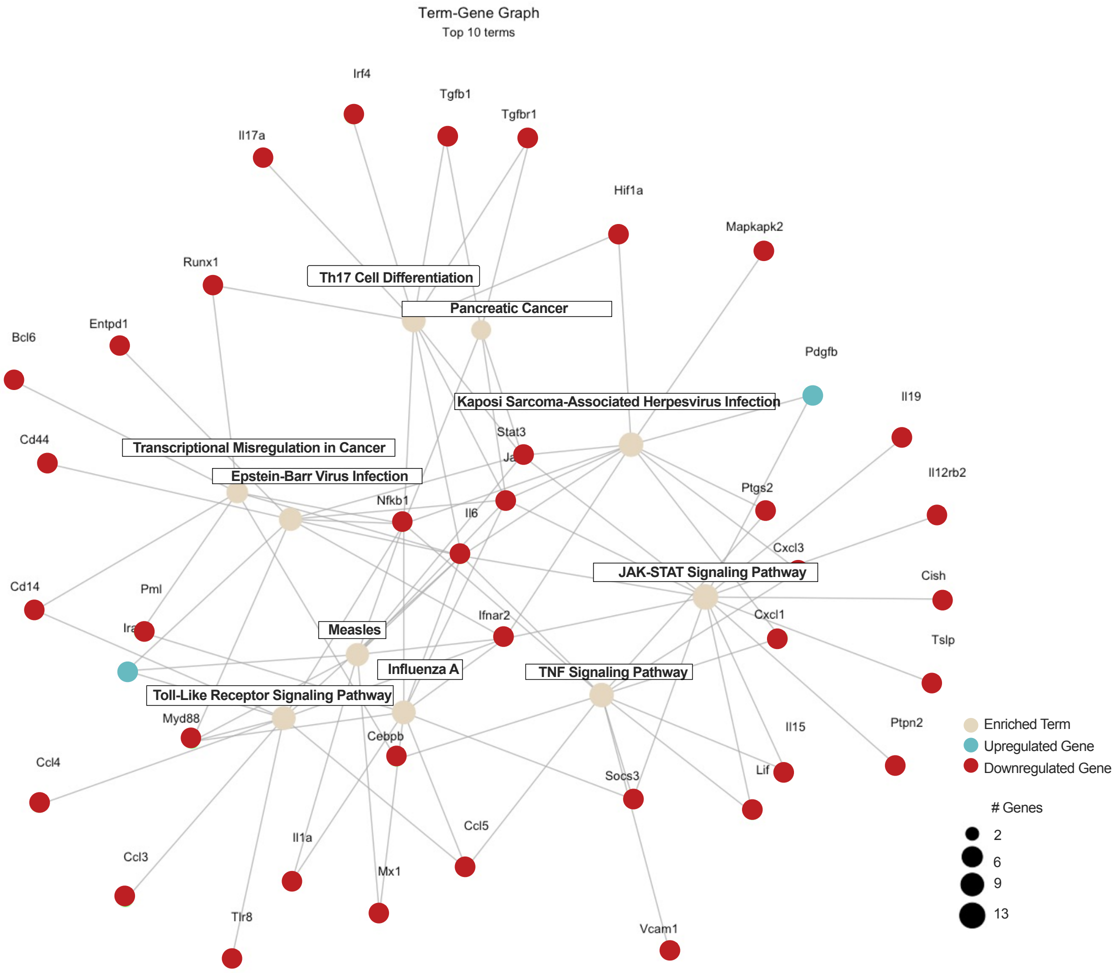


**Supplemental Figure 11. Gene map of significantly differentially expressed genes ($p < 0.05$) in lung samples in iECM vs saline-treated animals.** A total of n = 6 lung RNA samples were utilized and run on the Nanostring nCounter panel. Differential expression was determined using ROSALIND. Differentially expressed genes were inputted into R and analyzed using pathfindR. For visualization, only the top 5 differential pathways were used in this figure. Size of enriched term node is correlated to number of differentially expressed genes within that pathway. Genes upregulated with iECM treatment are shown in blue, while downregulated genes are visualized in red.

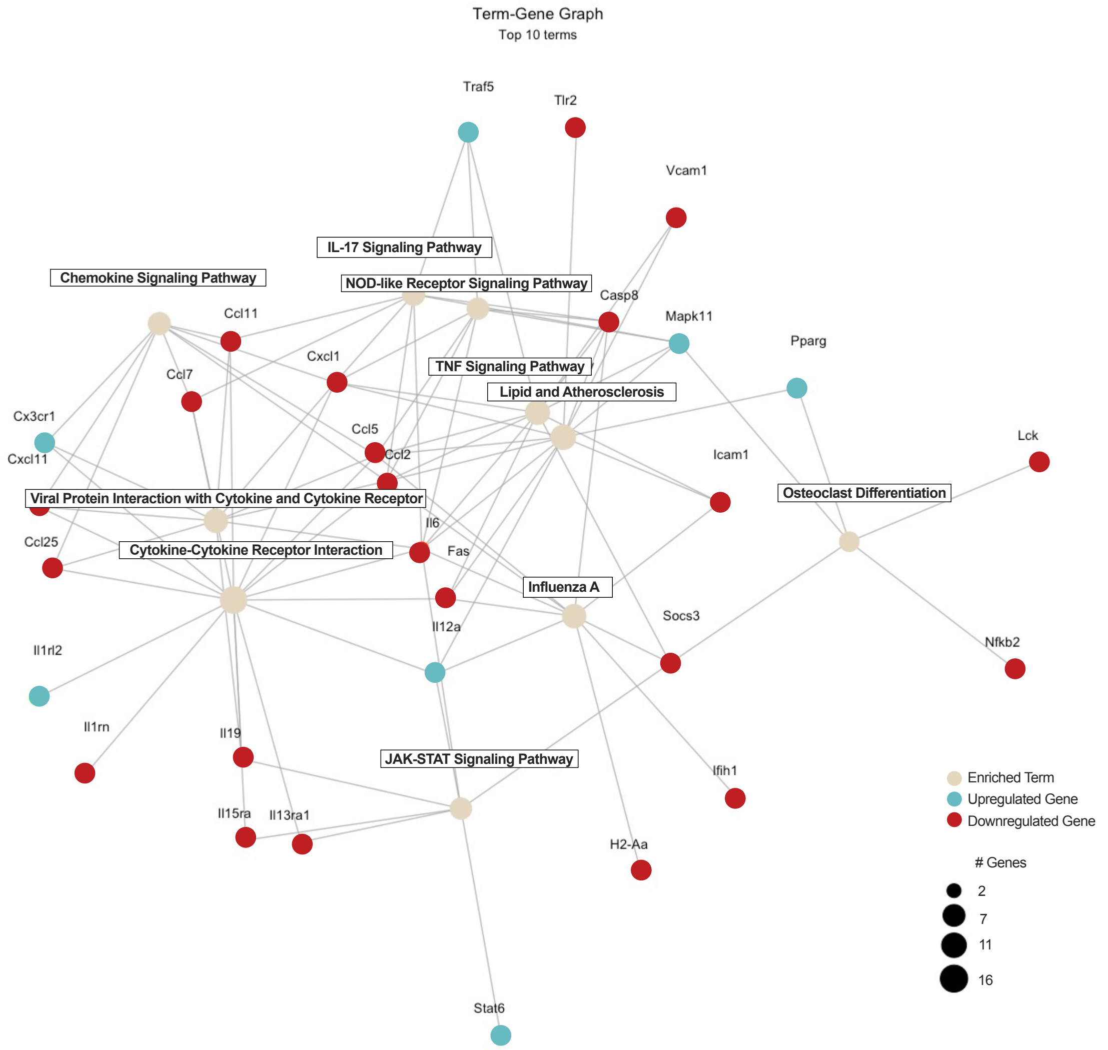


**Supplemental Figure 12. Gene map of significantly differentially expressed genes ($p < 0.05$) in brain samples in iECM vs saline-treated animals.** A total of n = 6 brain RNA samples were utilized and runu on the Nanostring nCounter panel. Differential expression was determined using ROSALIND. Differentially expressed genes were input into R and analyzed using pathfindR. Size of enriched term node is correlated to number of differentially expressed genes within that pathway. Genes upregulated with iECM treatment are shown in blue, while downregulated genes are visualized in red.

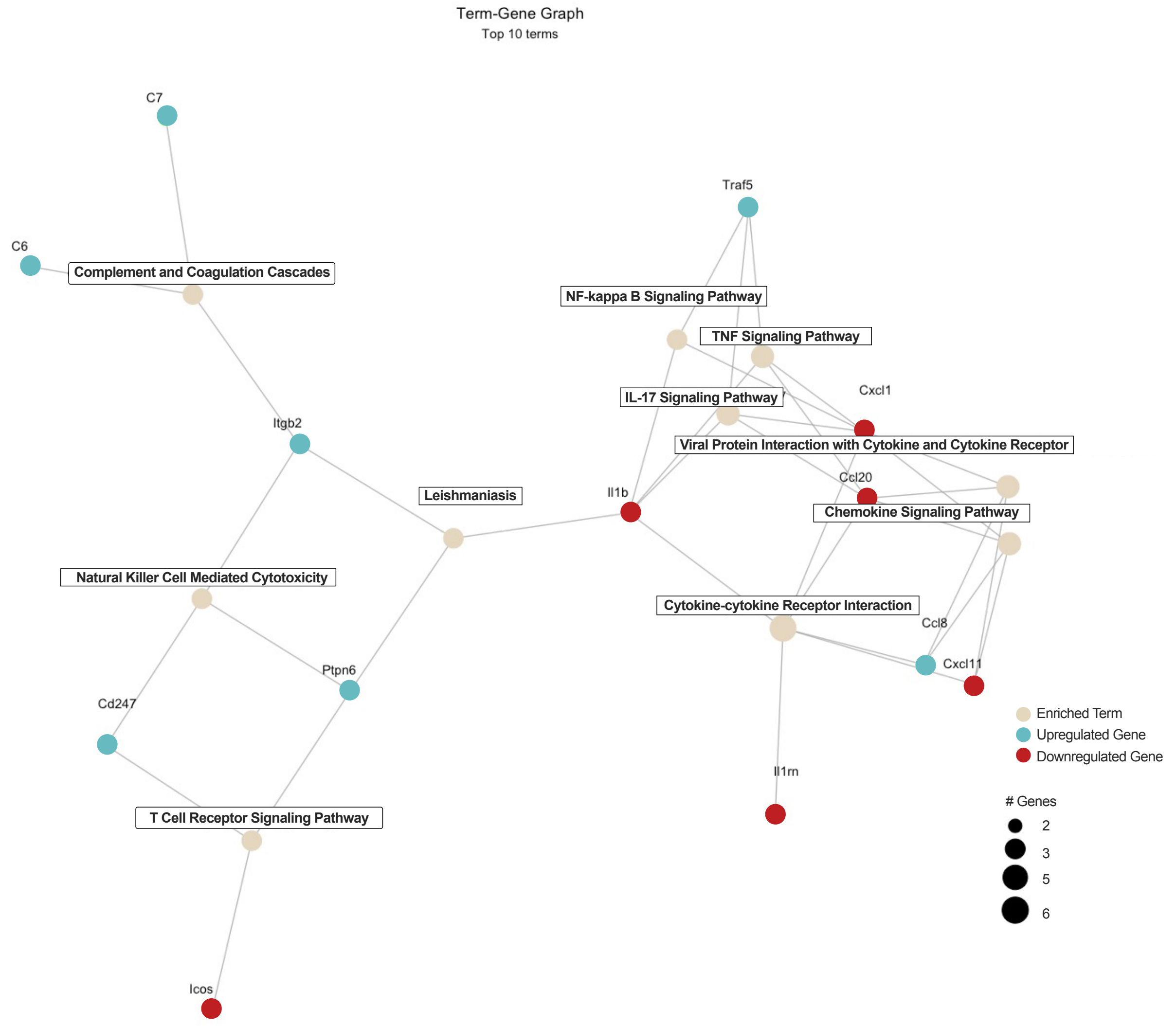


**Supplemental Figure 13. Gene map of significantly differentially expressed genes ($p < 0.05$) in kidney samples in iECM vs saline-treated animals.** A total of n = 6 kidney RNA samples were utilized and run on the Nanostring nCounter panel. Differential expression was determined using ROSALIND. Differentially expressed genes were inputted into R and analyzed using pathfindR. Size of enriched term node is correlated to number of differentially expressed genes within that pathway. Genes upregulated with iECM treatment are shown in blue, while downregulated genes are visualized in red.

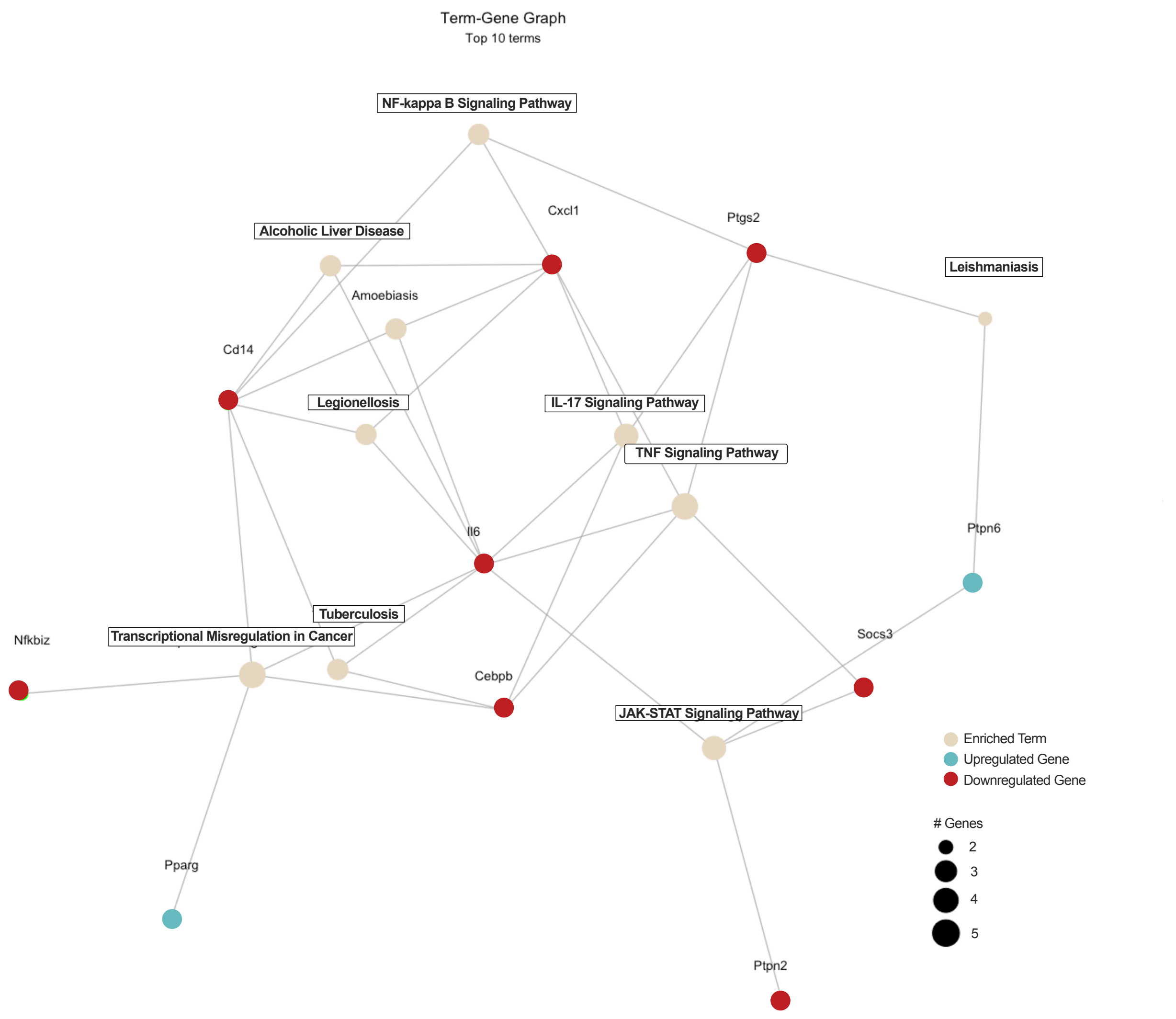


**Supplemental Figure 14. Gene map of significantly differentially expressed genes ($p < 0.05$) in heart samples in iECM vs saline-treated animals**. A total of n = 6 heart RNA samples were utilized and run on the Nanostring nCounter panel. Differential expression was determined using ROSALIND. Differentially expressed genes were inputted into R and analyzed using pathfindR. Size of enriched term node is correlated to number of differentially expressed genes within that pathway. Genes upregulated with iECM treatment are shown in blue, while downregulated genes are visualized in red.

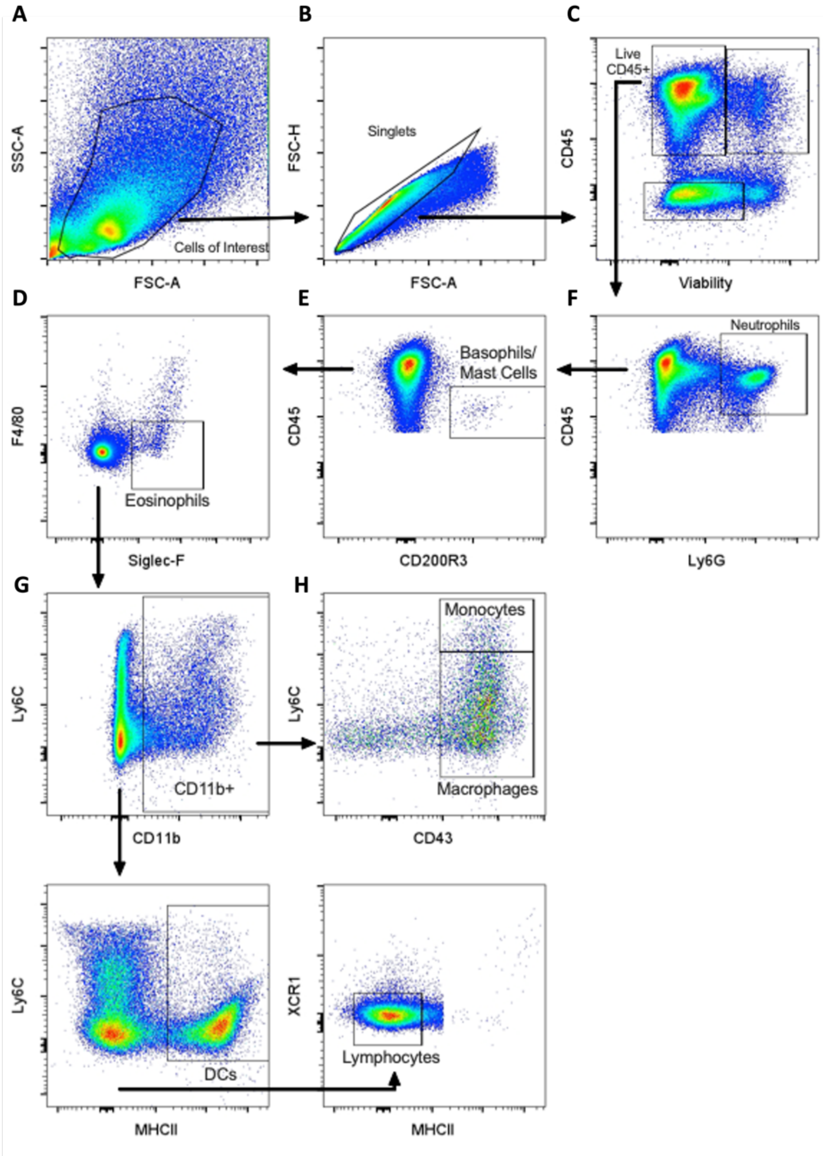


**Supplemental Figure 15. Gating strategy for 22 color myeloid panel in lung samples of LPS-dosed animals.** (**a**) Cells were first gated by forward and side scatter for correct size and granularity. (**b**) Singlets were isolated from doublets and cell clusters. (**c**) Live CD45+ cells were gated on based on low viability dye positivity and CD45 positivity. (**d**) Neutrophils were determined as Ly6G+ cells from CD45+ live cells. (**e**) Basophils/mast cells were identified by CD200R3 positivity from the non-neutrophil population. (**f**) Eosinophils were gated on from non-basophils/mast cells based on Siglec-F positivity and F4/80 negativity to eliminate alveolar macrophages. (**g**) Non-eosinophils were separated based on CD11b expression. (**h**) From CD11b+ cells, monocytes and macrophages were identified based on Ly6C and CD43 expression. Monocytes were determined as CD43+, Ly6Chi, while macrophages were identified by CD43 expression. (**i**) From CD11blo and CD11b- populations, dendritic cells were identified based on expression of MHCII. (**j**) Non-MHCII+ cells were isolated again and determined as lymphoid cells.

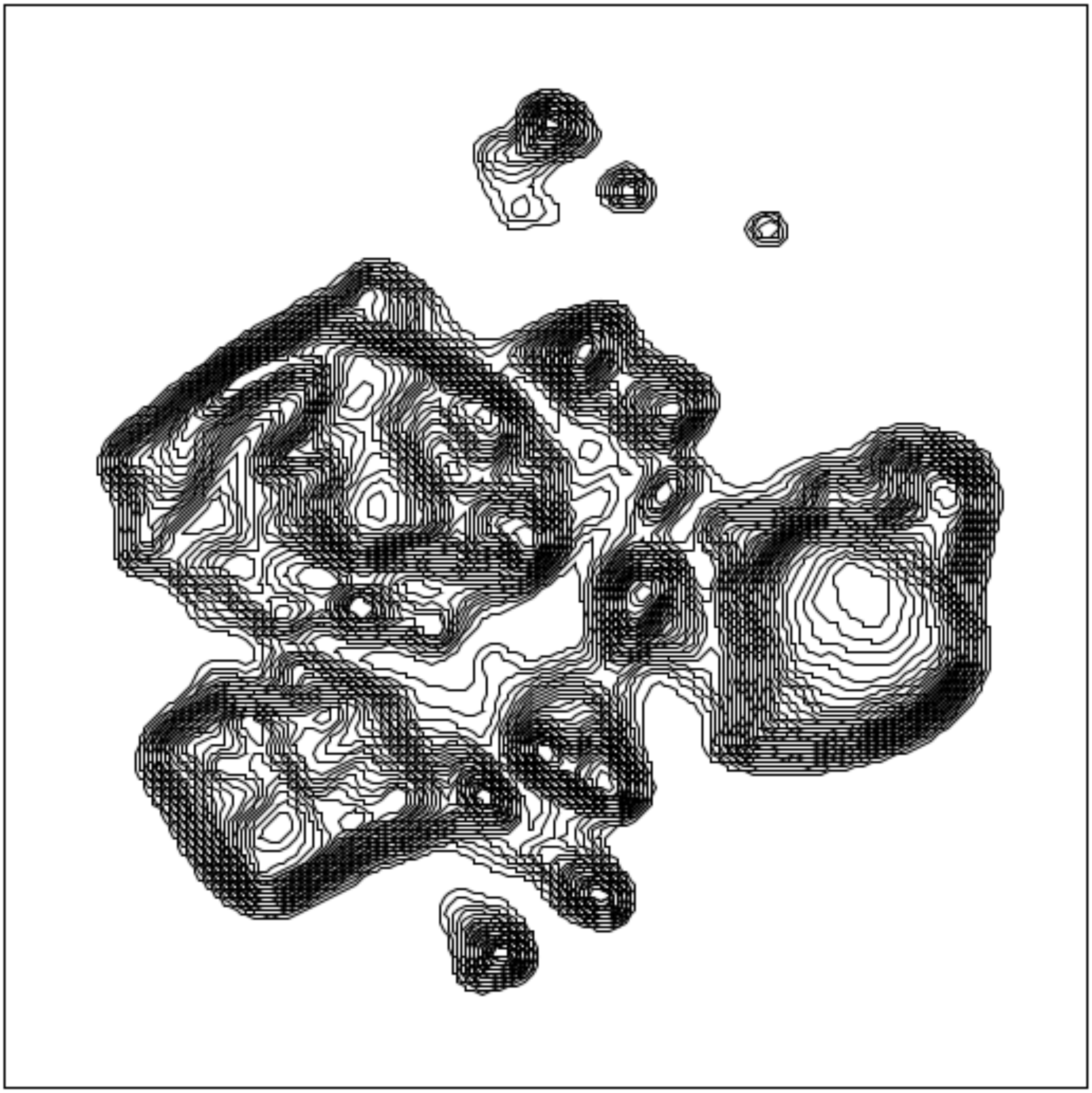

**Supplementary Figure 16. UMAP of all live CD45+ cells from concatenated iECM and saline samples.** Live CD45+ cells were determined in flow cytometry samples using gating on CD45 positive, Live/Dead Blue negative cells.

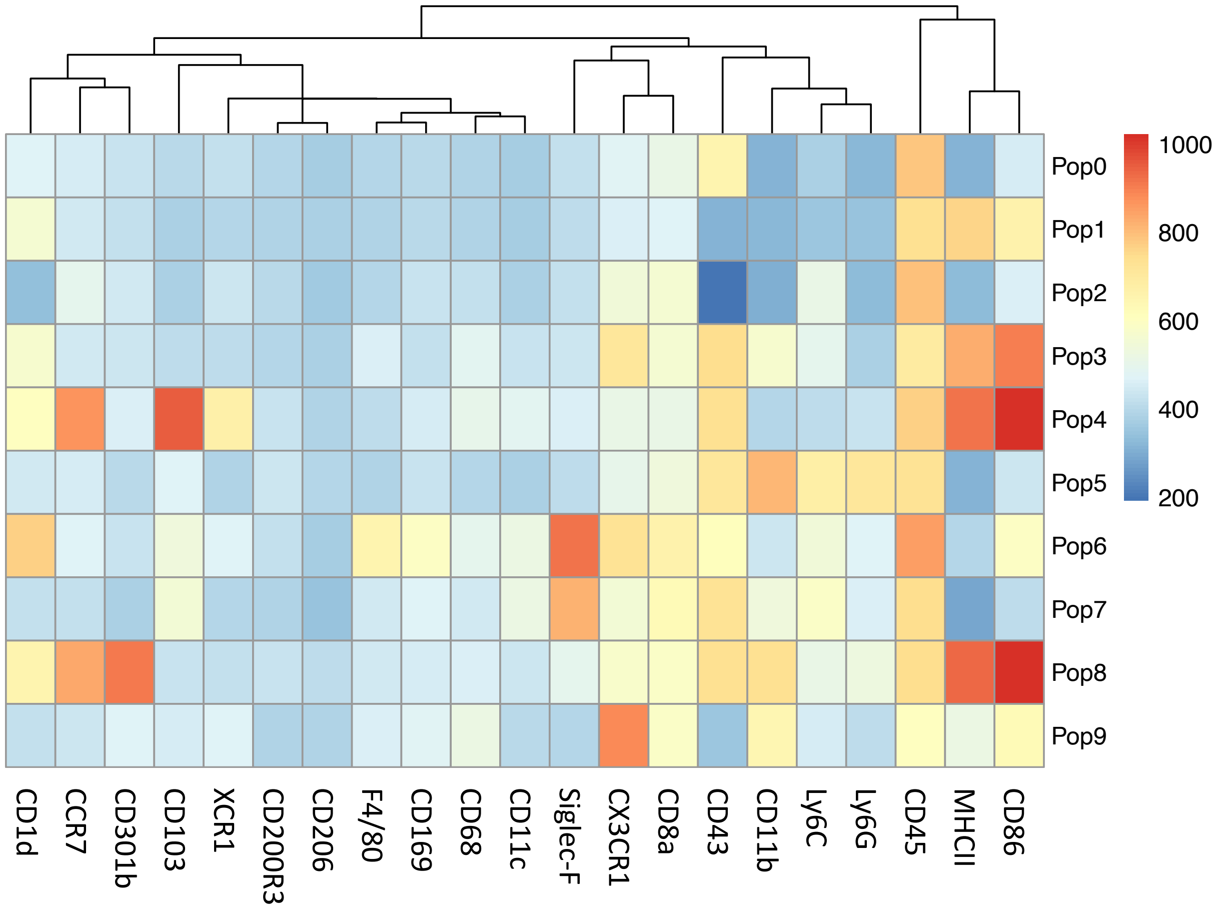


**Supplemental Figure 17. Heat map of clusters from unsupervised clustering of CD45+ live cells in LPS-dosed, saline and iECM-treated animals using FlowSOM.**

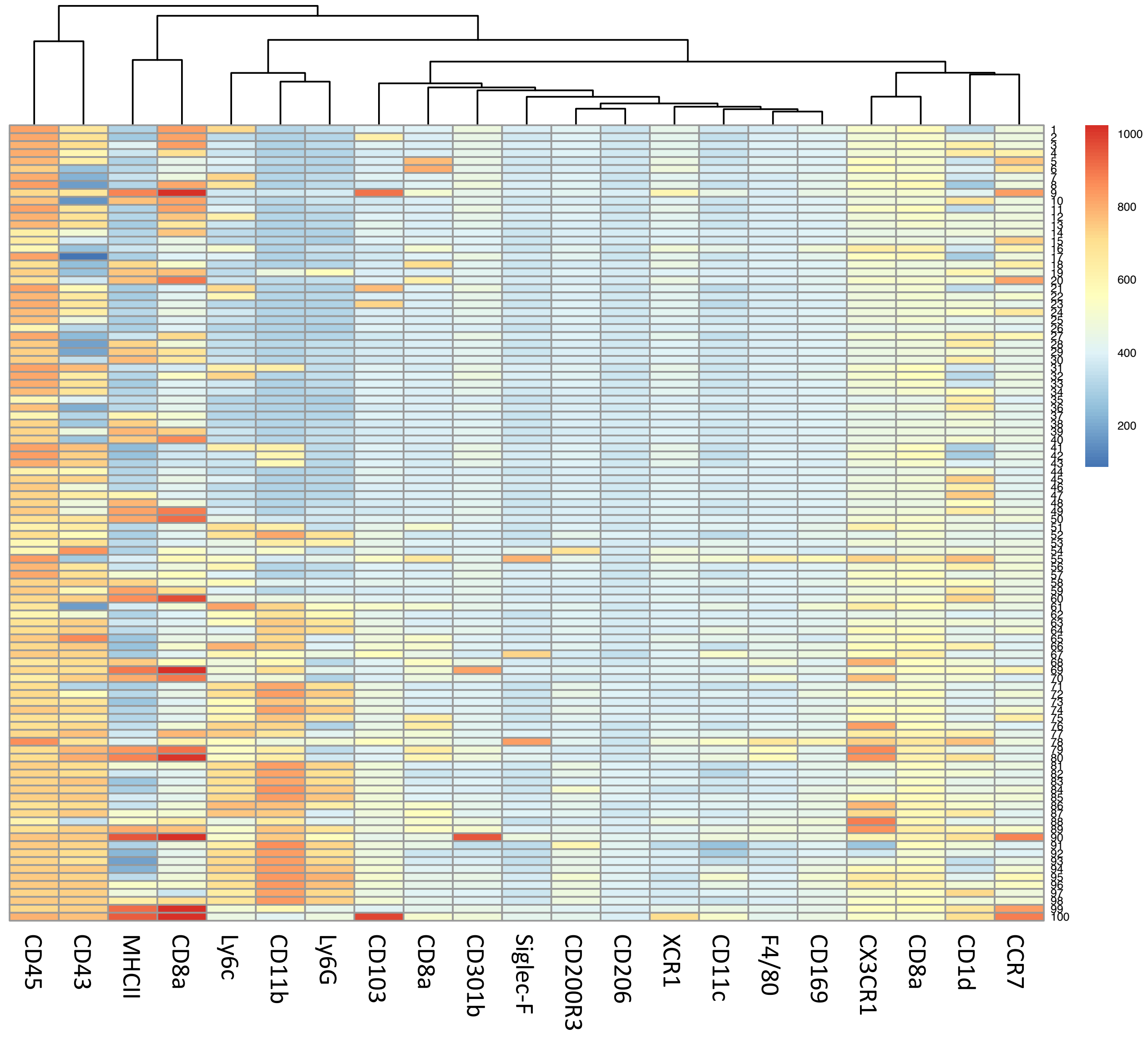


**Supplemental Figure 18**: **Heatmap of subclusters from unsupervised clustering of CD45+ live cells in LPS-dosed, saline and iECM-treated animals using flowSOM.**

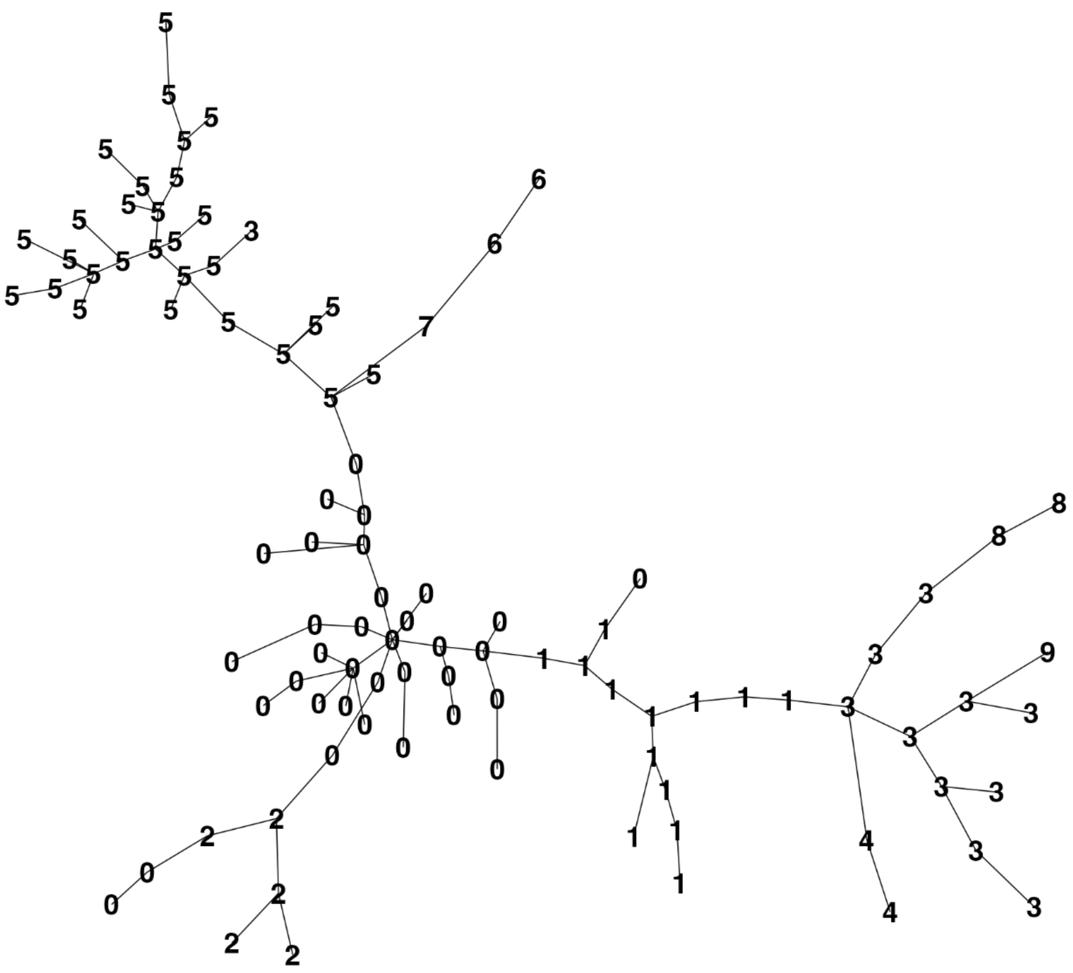

**Supplemental Figure 19. Hierarchy tree map of FlowSOM-developed unsupervised clustering of live CD45+ cell populations in saline and iECM-treated animals.**

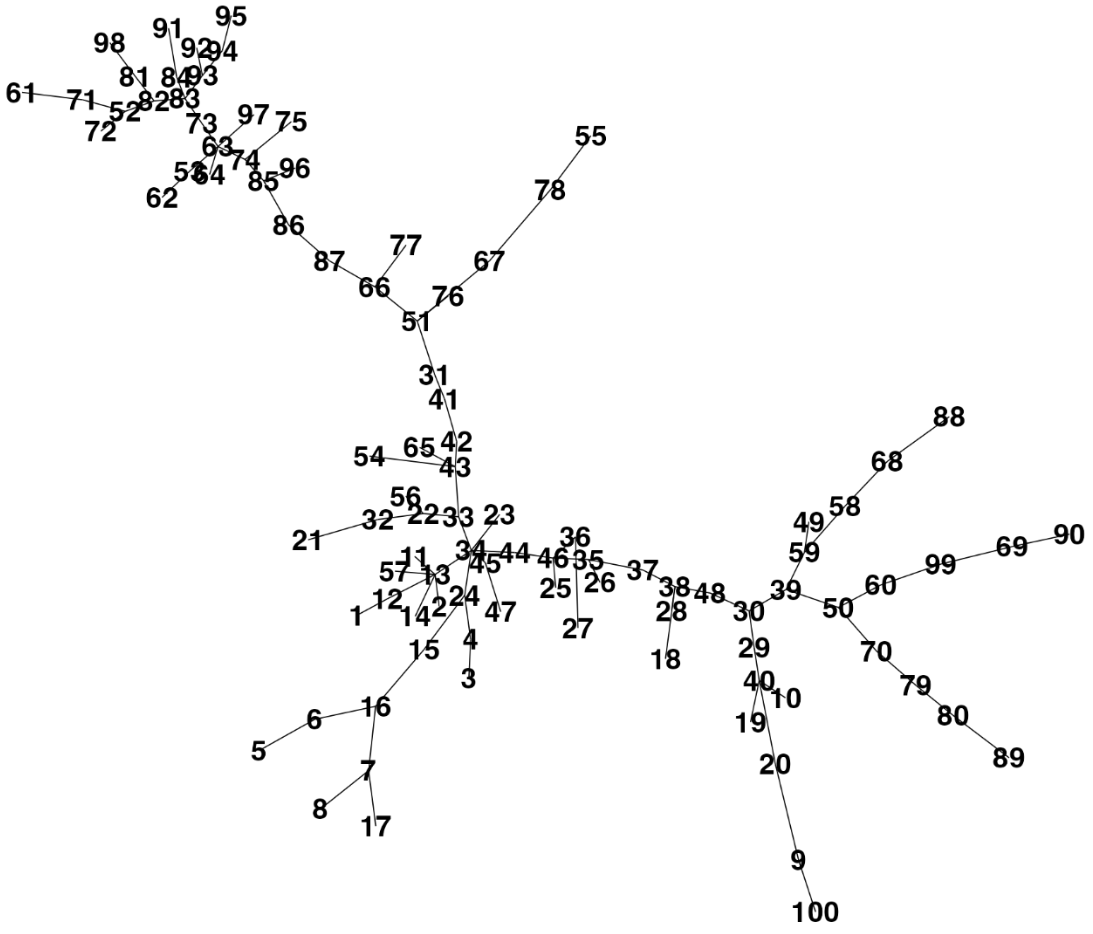


**Supplemental Figure 20. Subclusters of live CD45+ cells in LPS-dosed, saline and iECM-treated animals using FlowSOM unsupervised clustering.**

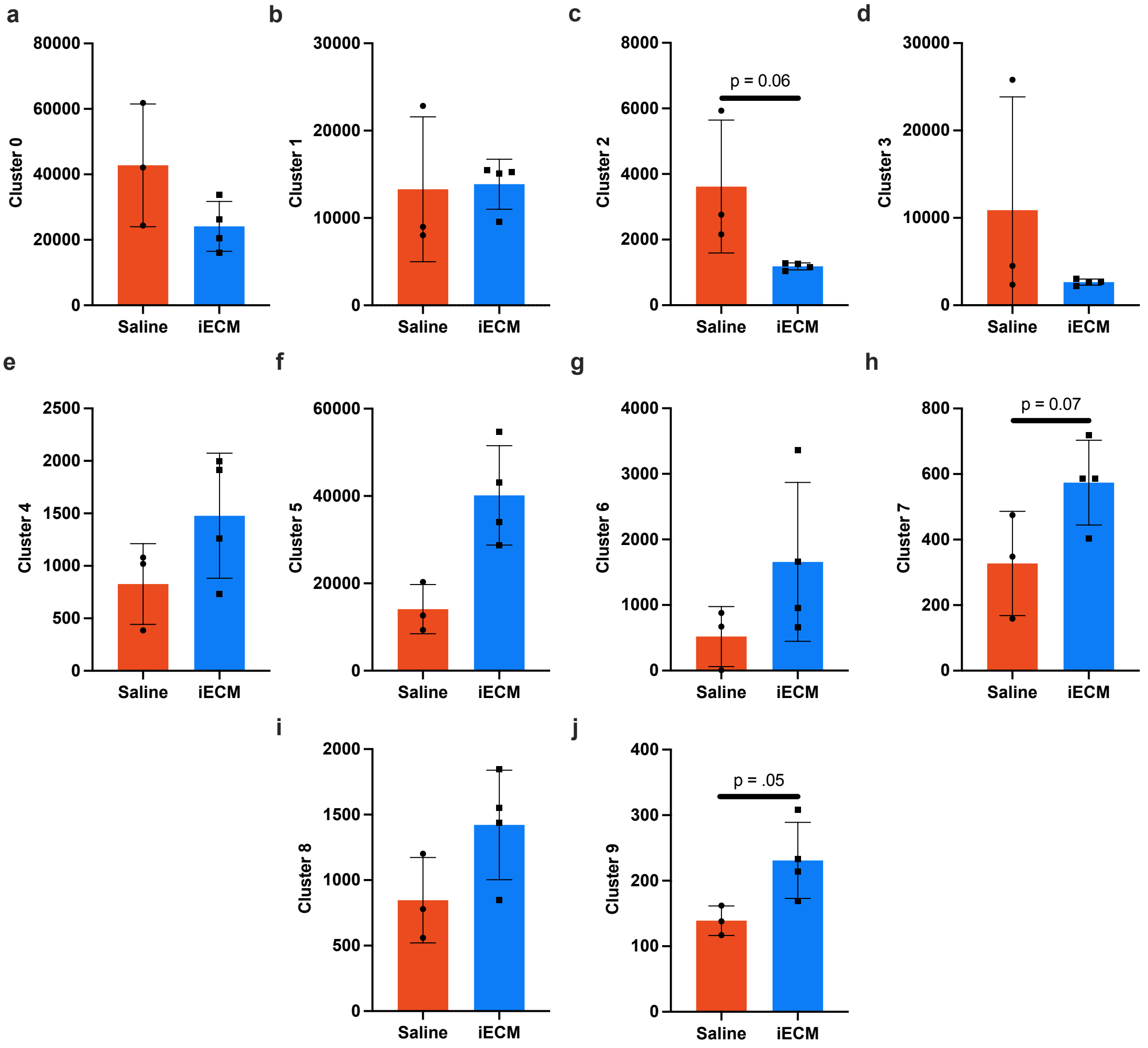


**Supplementary Figure 21. Levels of flowSOM cluster (a) 0 (b) 1 (c) 2 (d) 3 (e) 4 (f) 5 (g) 6 (h) 7 (i) 8 and (j) 9 in saline (red) and iECM- (blue) treated animals.** Cluster population levels are portrayed as number of cells as concatenated samples where equal counts of cells were taken from each sample. Significance was determine utilizing an unpaired t-test where significant different were determined as $p < 0.05$. Data is mean ± standard deviation.

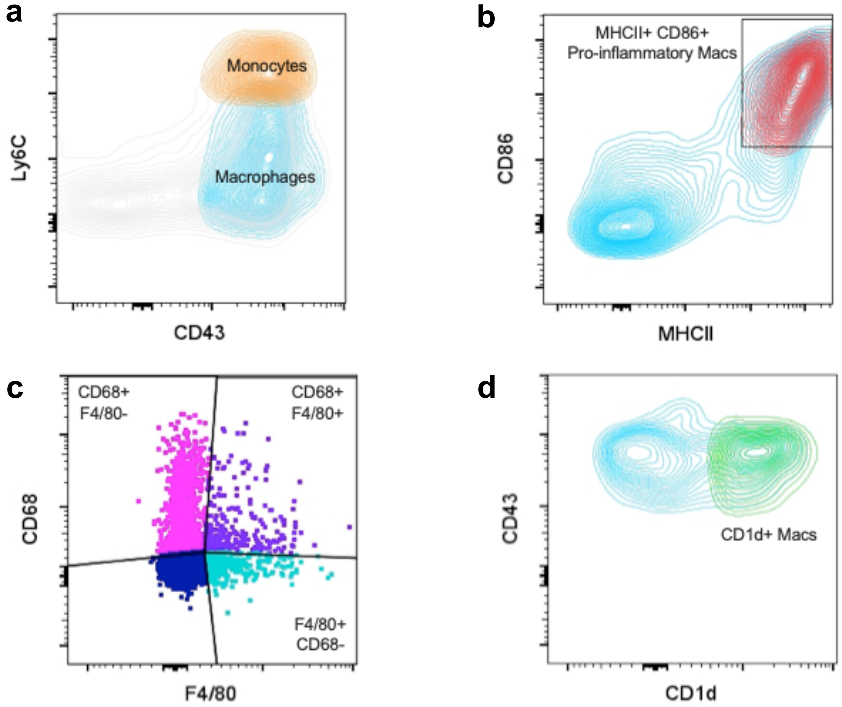


**Supplemental Figure 22. Identification of macrophage phenotypes using spectral flow cytometry.** **(a)** Macrophages were isolated from CD45+, CD11b+ cells expressing CD43 and low to medium levels of Ly6C. **(b)** MHCII and CD86 positivity were used to identify pro-inflammatory 'M1' type macrophages. **(c)** Macrophages were subclustered based on CD68 and F4/80 expression. **(d)** CD1d expression gating strategy for macrophages isolated CD1d+ macrophages.

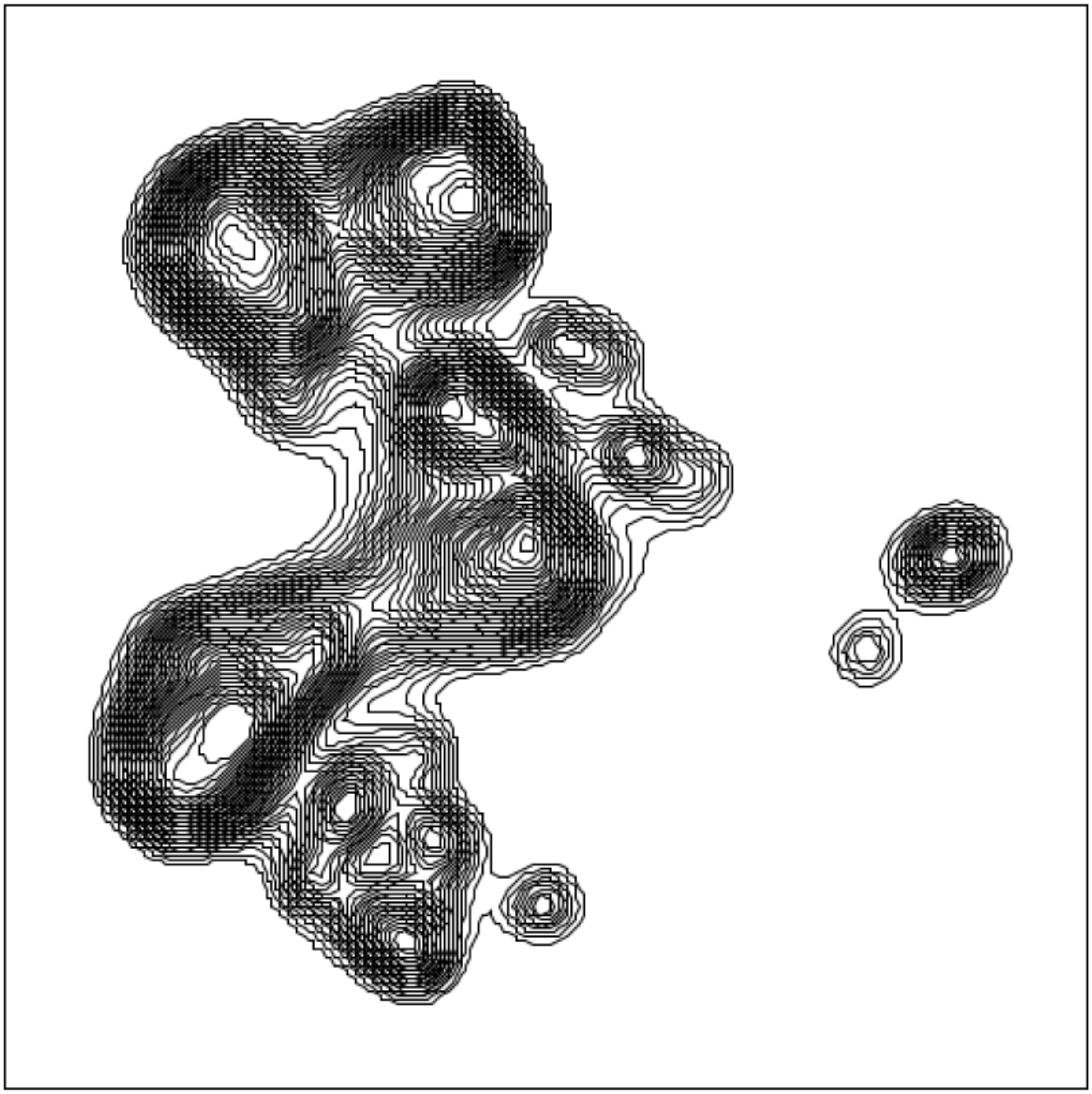

**Supplementary Figure 23. UMAP of macrophages in saline and iECM-treated animals.**

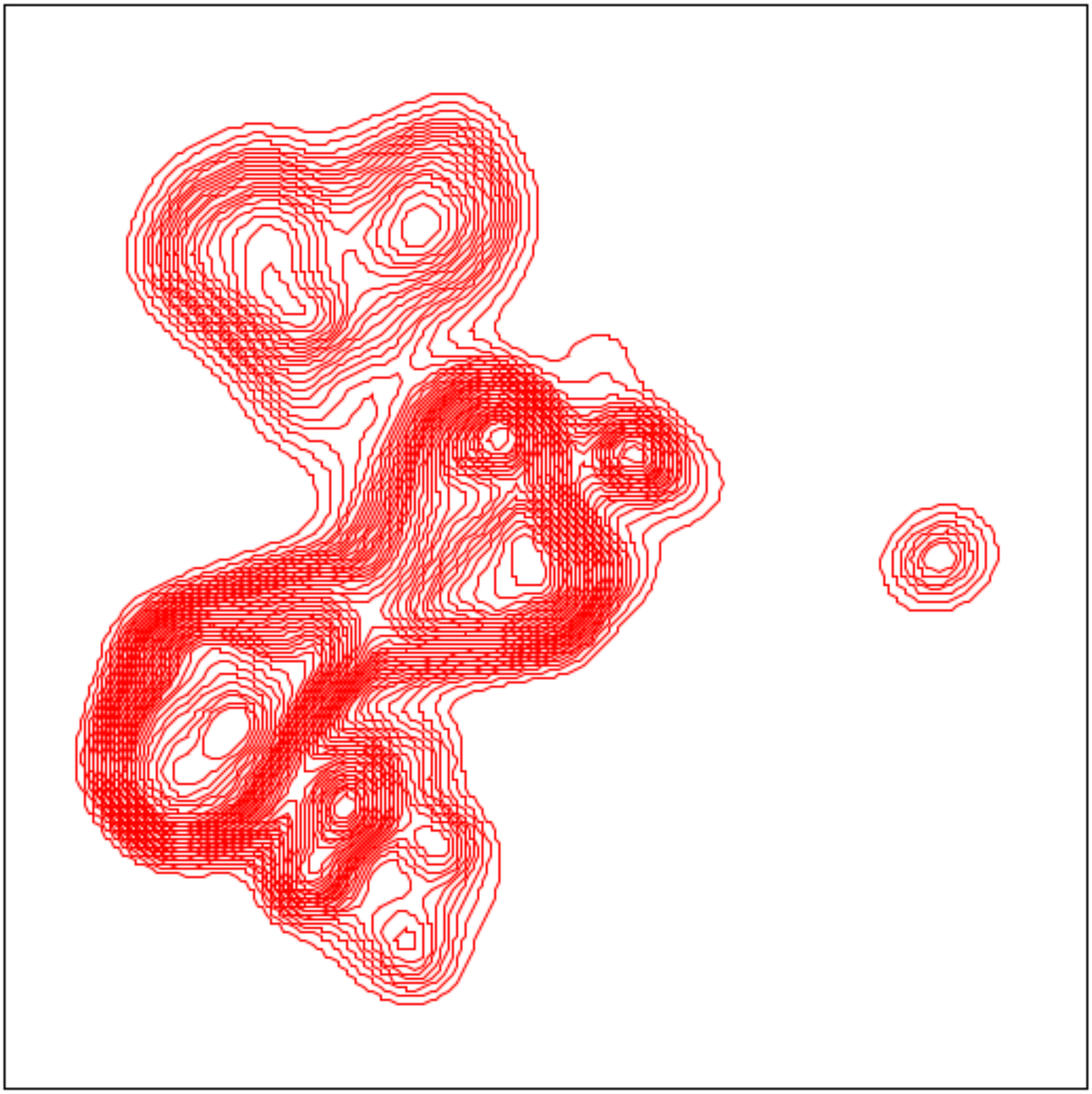

**Supplementary Figure 24. UMAP of macrophages in saline-treated animals.**

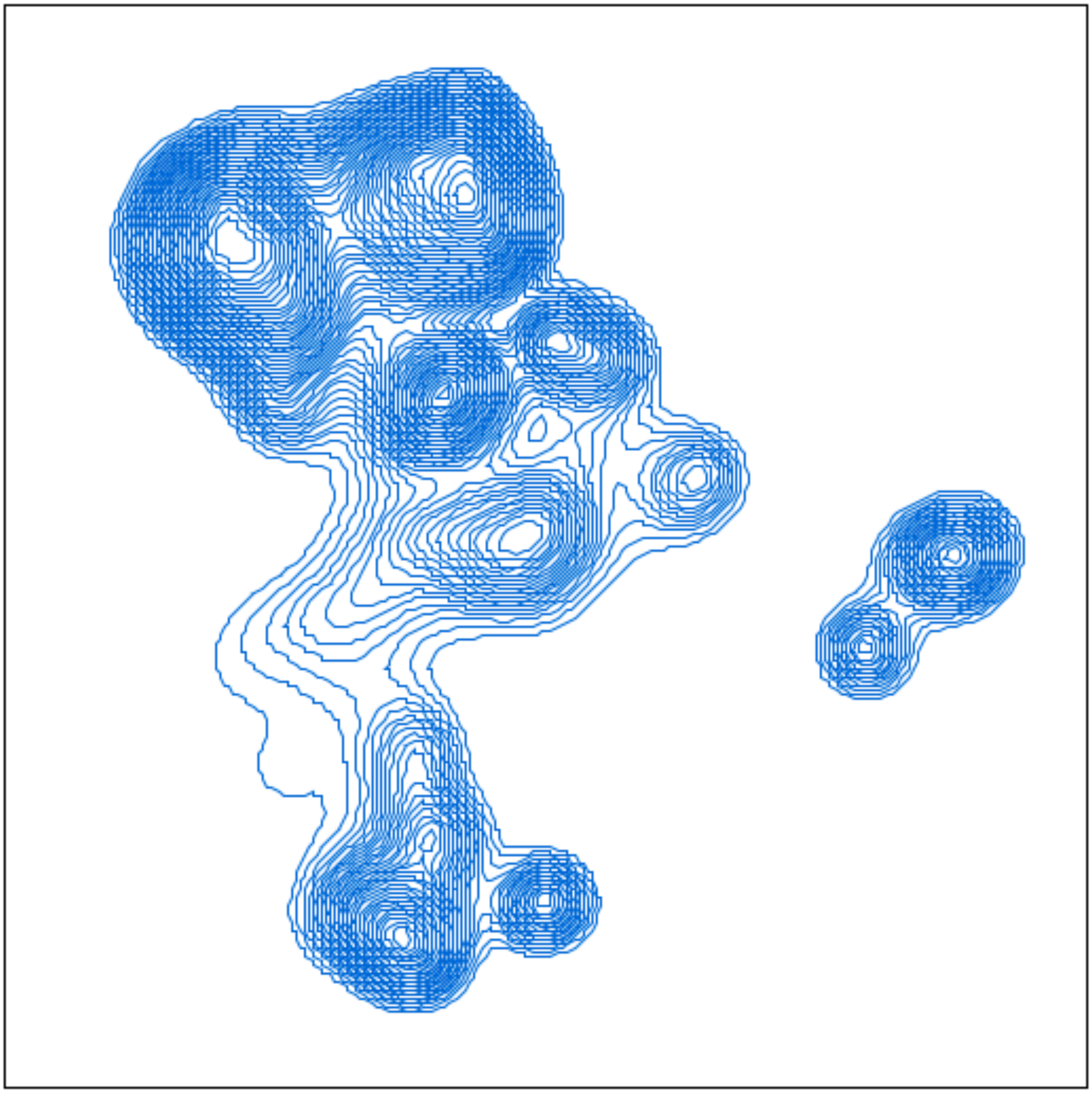

**Supplementary Figure 25. UMAP of macrophages in iECM-treated animals.**

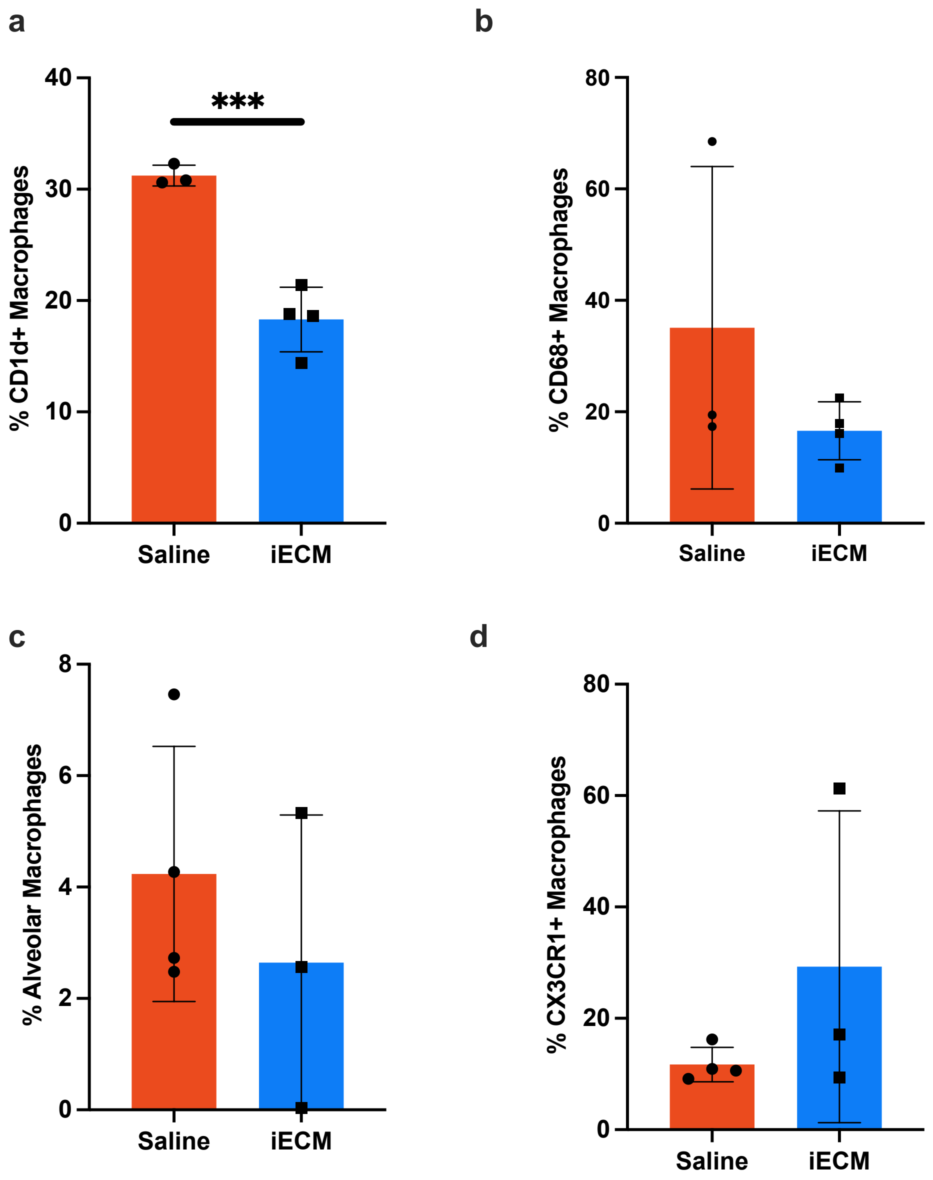


**Supplementary Figure 26. Macrophage characterization in saline versus iECM-treated animals.** A. CD1d macrophage expression was determined by CD1d expression in macrophages as seen in Supp. Figure 17. B. CD68 activated macrophage levels were determined based on CD68 expression. C. The percentage of alveolar macrophages were determined based on expression of SiglecF on macrophages. D. Levels of CX3CR1+ macrophages were assessed based on CX3CR1 expression as shown in Supp. Figure 17. Data is mean ± standard deviation. Significance was determined using a two-tailed unpaired t-test. p***<0.001

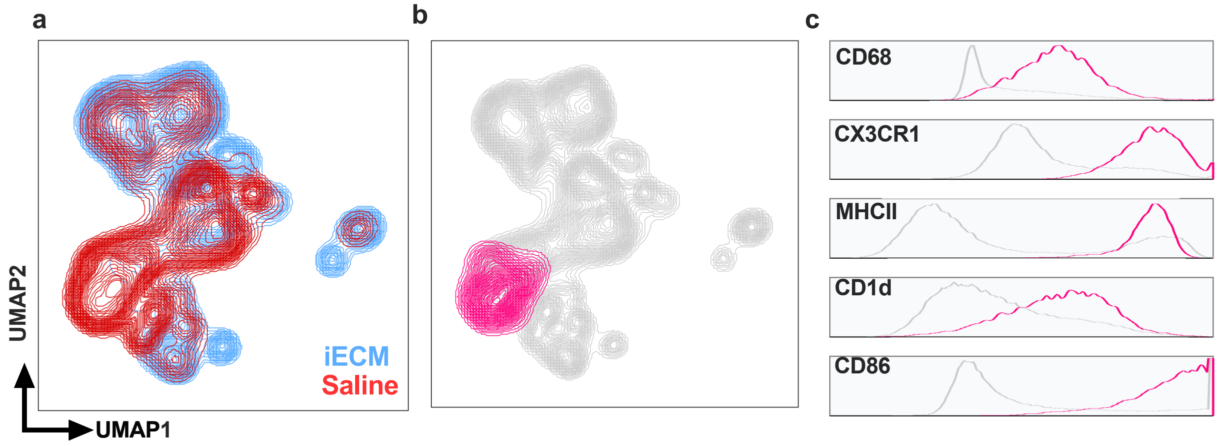


**Supplementary Figure 27.** Analysis of unique macrophage cluster in lungs of saline-treated animals. A. Macrophages were determined using the gating strategy outlined in Supplementary Figure 8. Unbiased clustering of macrophage subpopulations was performed using the UMAP plugin in FlowJo. B. Isolation of a saline-unique population of macrophages (pink) relative to other macrophage populations (gray) C. Histogram characterization of pink macrophage cluster unique to saline-treated animals for CD68, CX3CR1, MHCII, CD1d, and CD86 expression relative to all other macrophage populations in gray.
